# Asymbiotic mass production of the arbuscular mycorrhizal fungus *Rhizophagus clarus*

**DOI:** 10.1101/2020.12.25.424379

**Authors:** Sachiko Tanaka, Kayo Hashimoto, Yuuki Kobayashi, Koji Yano, Taro Maeda, Hiromu Kameoka, Tatsuhiro Ezawa, Katsuharu Saito, Kohki Akiyama, Masayoshi Kawaguchi

**Affiliations:** Division of Symbiotic Systems, National Institute for Basic Biology, 38 Nishigonaka, Myodaiji, Okazaki, Aichi 444-8585, Japan; Graduate School of Life and Environmental Sciences, Osaka Prefecture University, 1-1 Gakuen-cho, Nakaku, Sakai, Osaka 599-8531, Japan; Graduate School of Agriculture, Hokkaido University, Kita 9, Nishi 9, Kita-ku, Sapporo, Hokkaido 060-8589, Japan; Faculty of Agriculture, Shinshu University, 8304 Minamiminowa, Nagano 399-4598, Japan; Department of Basic Biology, School of Life Science, Graduate University for Advanced Studies (SOKENDAI), 38 Nishigonaka, Myodaiji, Okazaki 444-8585, Japan

## Abstract

Arbuscular mycorrhizal (AM) symbiosis is a mutually beneficial interaction between fungi and land plants and promotes global phosphate cycling in terrestrial ecosystems. AM fungi are recognised as obligate symbionts that require root colonisation to complete a life cycle involving the production of propagules, asexual spores. Recently it has been shown that *Rhizophagus irregularis* can produce infection-competent secondary spores asymbiotically by adding a fatty acid, palmitoleic acid. Further, asymbiotic growth can be supported using myristate as a carbon and energy source for their asymbiotic growth to increase fungal biomass. However, spore production and the ability of these spores to colonise host roots were still limited compared to co-culture of the fungus with plant roots. Here we show that a combination of two plant hormones, strigolactone and methyl jasmonate, induces production of a large number of infection-competent spores in asymbiotic cultures of *Rhizophagus clarus* HR1 in the presence of myristate and organic nitrogen. Inoculation of asymbiotically-generated spores promoted the growth of Welsh onions, as observed for spores produced by symbiotic culture system. Our findings provide a foundation for elucidation of hormonal control of the fungal life cycle and development of new inoculum production schemes.

## Main

Arbuscular mycorrhizal (AM) fungi are ubiquitous symbionts of the majority of terrestrial plant species and can facilitate the plant mineral acquisition^1,2^. AM fungi are obligate biotrophic fungi, depending on host-derived substrates including carbohydrates, such as sugars and lipids^3–6^. Genome analyses of AM fungi demonstrate that they lack several important metabolic enzymes related to the obligate biotrophy^7–11^. AM fungi have long been considered unculturable without the host. However, co-culture of the AM fungus *Rhizophagus irregularis* and mycorrhiza-helper bacteria–isolates of *Paenibacillus validus–demonstrated* that AM fungi can complete their life cycle in the absence of host plants^12,13^. Recently, fatty acids have been shown to boost AM fungal growth and sporulation under asymbiotic conditions. Palmitoleic acid asymbiotically induced infection-competent secondary spores of *R. irregularis*^14^. Further, myristate initiated the asymbiotic growth of AM fungi and can also serve as a carbon and energy source^15^. These findings may lead to the development of new research tools for AM studies and novel generation system of AM fungal inoculants. However, at present, fungal biomass and spore production in the asymbiotic culture systems remain lower than those in symbiotic co-cultures. Moreover, spores induced by palmitoleic acid or myristate were smaller than those generated symbiotically and their performance as inoculants are unknown.

Both nutrients and signalling molecules from host plants may be crucial for AM fungal growth and reproduction^16–18^. Some phytohormones show positive effects on interactions between AM fungi and hosts. Strigolactone is a major plant-derived signal known to induce hyphal branching^19^ and elongation^20^ of AM fungi and to stimulate their mitochondrial activity^21^ in the pre-symbiotic stage. Methyl jasmonate (MeJA) was increased during AM fungal colonization in roots^22^, consistent with the up-regulation of jasmonic acid biosynthesis genes in plant cortical cells containing arbuscules which are highly branched fungal structures for nutrient exchange^23,24^. Most research of these and other phytohormones are focused on cell-level interactions between AM fungi and plants, and the direct effect of phytohormones on AM fungal growth and reproduction is largely unclear.

Here, we focused on the effect of two phytohormones, strigolactone and MeJA, on AM fungal growth and sporulation in asymbiotic culture supplemented with potassium myristate and organic nitrogen. We found that the asymbiotic growth was superior to *R. irregularis* in *R. clarus* HR1, whose genome has been sequenced^7^. Finally, hundreds of times more secondary spores were produced in medium containing the two phytohormones than in media without phytohormones in *R clarus* asymbiotic culture. Furthermore, we confirmed that colonisation by asymbiotically-generated spores facilitates the growth of Welsh onions.

## Results

### Survey of base media containing fatty acids and organic nitrogen sources for asymbiotic culture of *R. clarus* HR1

We developed a new base medium for asymbiotic culture of *R. clarus* HR1. According to genomic analysis of *R. clarus*, this fungus lacks metabolic pathways of disaccharide degradation and thiamine biosynthesis^7^. Thus, we changed the composition of the original modified M medium by adding glucose and more thiamine and reducing sucrose (Supplementary Table 1), which was used for all culture experiments in this report. First, we assessed four fatty acids on *R. clarus* asymbiotic culture: potassium myristate, potassium palmitoleate, potassium palmitate and 2-hydroxytetradecanoic acid (2OH-TDA, which can promote hyphal elongation of *Gigaspora* spp.^25^). Ten parent spores, seed fungus collected from *in vitro* monoxenic culture, were placed separately on a solid medium and incubated for 6 weeks. Only potassium myristate unlike other tested fatty acids activated hyphal growth of *R. clarus* (Fig. 1a and Supplementary Fig. 1). In asymbiotic culture supplemented with potassium myristate, AM fungus expanded its habitat by generating straight, thick hyphae (runner hyphae; RH) with small bunches of short branches. However, sporulation rate, the number of secondary spore-forming parent spores per germinated parent spores, was very low 6 weeks after incubation (WAI) in the three trials (Fig. 1b). Similarly, almost no parent spores formed secondary spores by the application of the other fatty acids.

**Figure 1.**
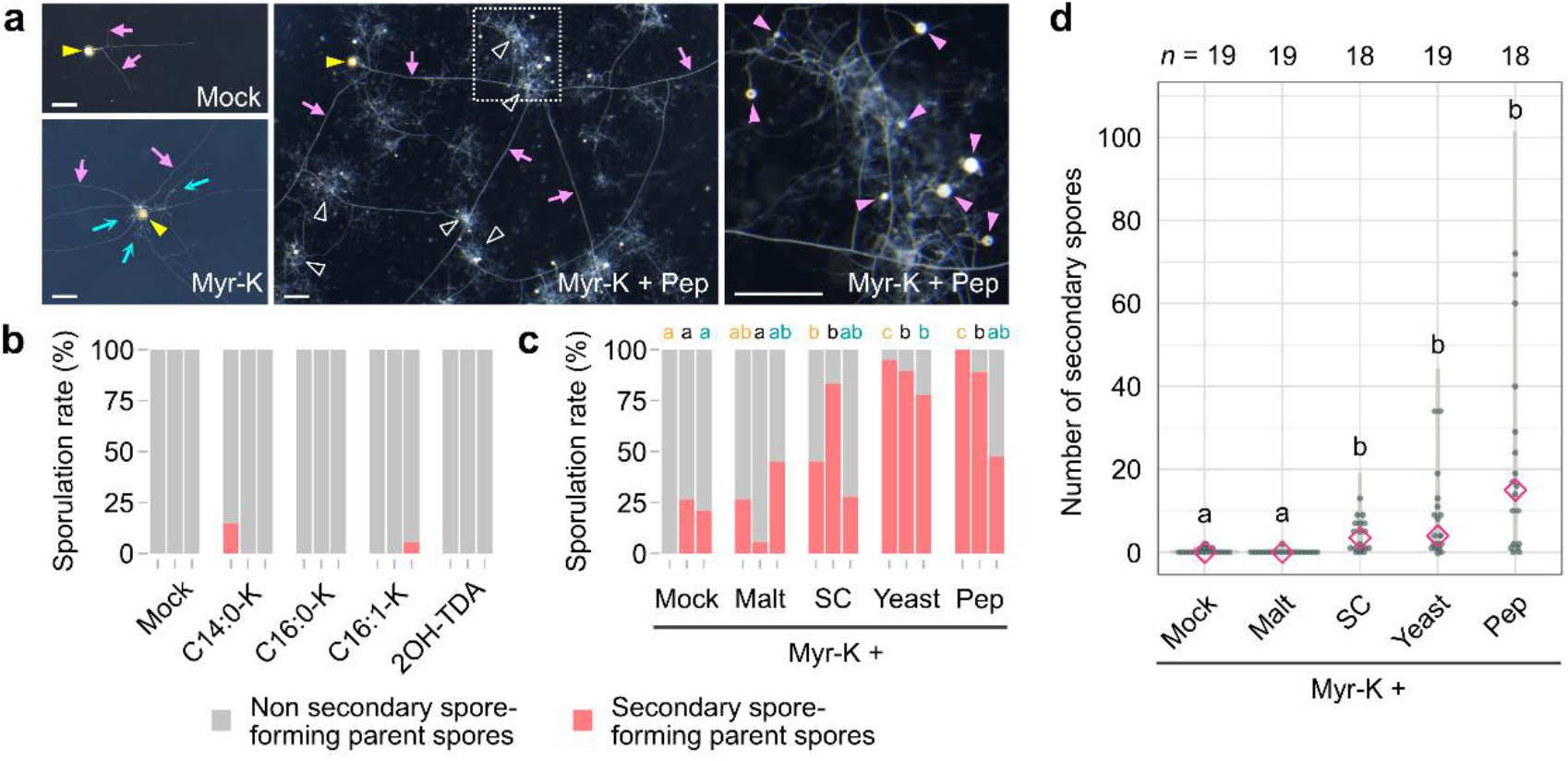
Effects of fatty acid salts and organic nitrogen sources on asymbiotic cultures of *R. clarus*. **a**, Effects of potassium myristate (Myr-K) and peptone (Pep) on *R. clarus* growth. AM fungi were incubated in medium with or without 100 μM Myr-K and 0.2 mg L^−1^ peptone for 6 weeks. Branched hyphae (blue arrows) were observed around a parent spore (yellow arrowhead) in medium containing Myr-K. Small, white spores were formed among densely packed coils (DPC, outlined arrowheads) in a medium containing both Myr-K and Pep. The rightmost picture is a magnified image of the dotted box in the middle picture. Pink arrowheads indicate newly generated secondary spores. Yellow arrows indicate runner hyphae. Bars = 500 μm. **b,** Sporulation rates, percentage of secondary spore-forming parent spores relative to germinated patent spores, in cultures supplemented with 100 μM fatty acid at 6 weeks after incubation (WAI). Three bars in each condition are trials 1 to 3, respectively from the left. 2OH-TDA, 2-hydroxytetradecanoic acid. C14:0-K, potassium myristate. C16:0-K, potassium palmitate. C16:1-K, potassium palmitoleate. **c**, Sporulation rates in cultures with 0.2 mg L^−1^ organic nitrogen source in the presence of Myr-K at 6 WAI. *Different letters* above of the graph indicate significant differences among treatments in each trial using Fisher’s exact test with Bonferroni correction (*p* < 0.05). Malt, malt extract. SC, SC dropout. Yeast, yeast extract. **d**, Numbers of secondary spores generated from a single parent spore in medium supplemented with each organic nitrogen in the presence of Myr-K at 6 WAI. Diamonds indicate medians. Statistical significance was calculated using the Wilcoxon rank-sum test with Bonferroni correction. *Different letters* indicate significant differences (*p* < 0.05). *p*-values are described in Supplementary Table 3.

In screening of substrates to induce secondary spore formation, we found that some organic nitrogen sources are effective for *R. clarus*. Fig. 1c shows the effect of four organic nitrogen sources (malt extract, SC dropout, yeast extract and peptone) on the sporulation in the presence of potassium myristate. All four organic nitrogen sources tended to enhance secondary spore formation, although sporulation rates varied among the three trials. In particular, the application of peptone showed high sporulation rate of 47 to 100% (Fig. 1c) and the highest production of secondary spores (Fig. 1d and Supplementary Fig. 2). *R. clarus* also produced many branched hyphae and densely packed coils (DPC)^12,13,15^, massive assemblies of thin hyphae, in the presence of peptone (Fig. 1a and Supplementary Fig. 2). No significant differences were found in secondary spore production in media supplemented with any combinations of 100 or 500 μM potassium myristate and 0.2 or 1.0 mg L^−1^ peptone (Supplementary Fig. 2). Taken together, we chose potassium myristate and peptone as components of the base medium for the following analyses of phytohormones in *R. clarus* asymbiotic culture. In this survey of base medium, we observed large variations in sporulation rate and number of secondary spores among trials. This is probably attributed to physiological status of each parent spore because spore maturity and degree of spore dormancy may differ among spores produced by *in vitro* monoxenic culture and the season when experiments were performed (Supplementary Table 2). Accordingly, we performed three independent cultures in the following experiments.

### Strigolactone enables the production of a greater number of secondary spores in asymbiotic culture

We examined the effect of the synthetic strigolactone GR24 on secondary spore formation in *R. clarus*. GR24 strongly stimulated fungal sporulation in the presence of potassium myristate and peptone, resulting in the generation of secondary spores in almost all germinated parent spores at 6 WAI (Fig. 2a). Notably, the median numbers of secondary spores in the presence of GR24 were increased by more than 10-fold compared with those in the mock treatment (Fig. 2b and Supplementary Fig. 3). We further analysed the early response of *R. clarus* to GR24. The application of GR24 accelerated germination of parent spores, which reached to more than 70% germination rate already at 5 days after incubation (DAI) (Fig. 2c). In contrast, spore germination in the absence of GR24 was less than 40% at 8 DAI and increased thereafter. Secondary spore formation was also accelerated by GR24 (Fig. 2d). Secondary spores emerged in culture media with GR24 by 8 DAI and the sporulation rate was increased to 80% at 14 DAI, whereas only a few germinated parent spores formed secondary spores during the first two weeks of culture without GR24. Thus, GR24 can promote germination of parent spores and secondary spore formation in *R. clarus*, which finally leads to almost 100% of sporulation rate and the production of a large number of secondary spores under asymbiotic conditions. We referred to a base medium containing both 500 μM potassium myristate and 1.0 mg L^−1^ peptone as T medium, and T medium containing 100 nM GR24 as TG medium.

**Figure 2.**
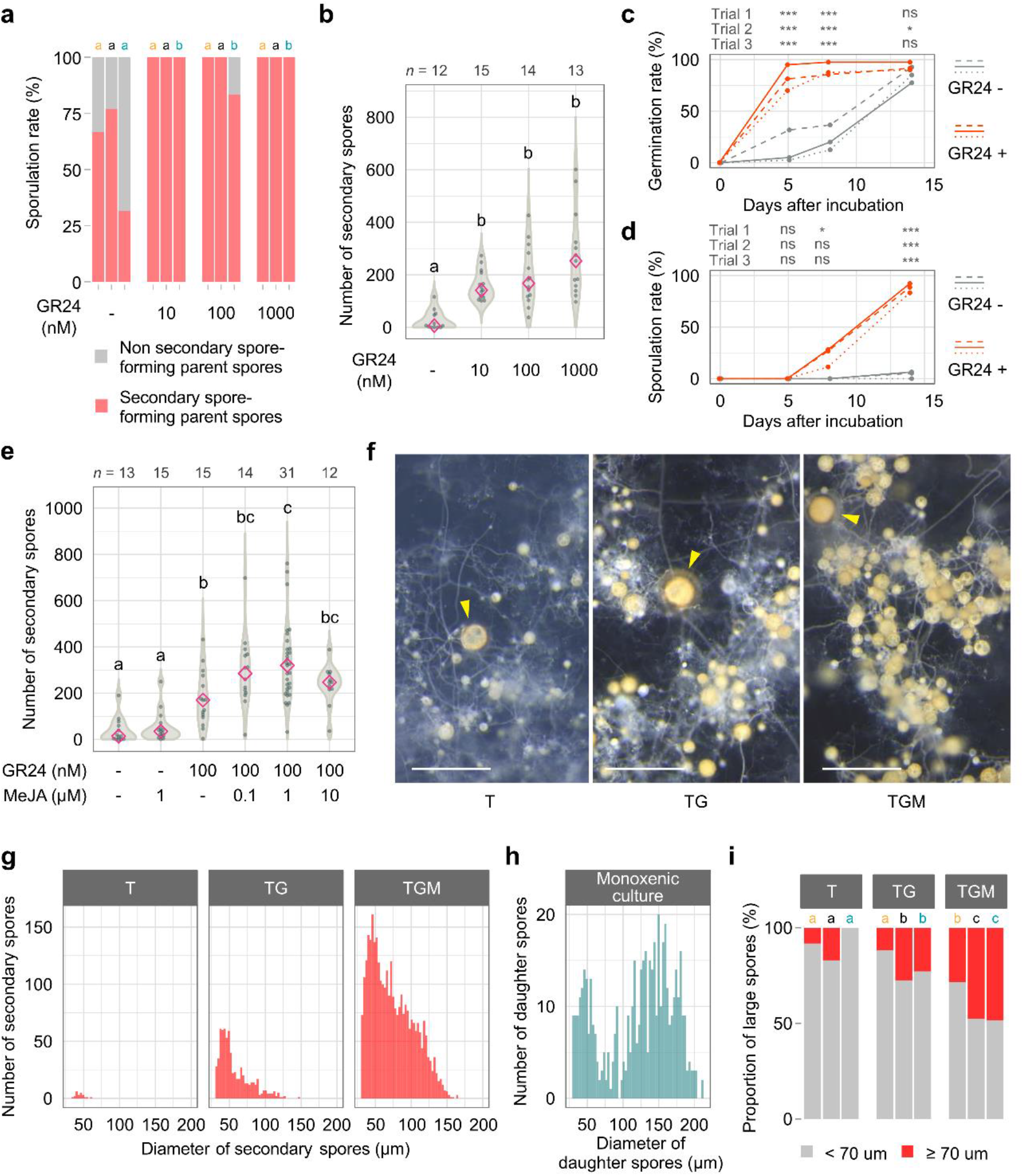
Effects of phytohormones on *R. clarus* asymbiotic cultures. All culture media contain both 500 μM Myr-K and 1 mg L^−1^ peptone. **a**, Percentage of secondary spore-forming parent spores relative to germinated patent spores in cultures with different concentrations of GR24 at 6 WAI. Three bars in each condition are trials 1 to 3, respectively from the left. *Different letters* above the graph indicate significant differences among treatments in each trial by Fisher’s exact test with Bonferroni correction (*p* < 0.05). **b**, Numbers of secondary spores in cultures with different concentrations of GR24 at 6 WAI. Diamonds indicate medians. Statistical significance was calculated using the Wilcoxon rank-sum test with Bonferroni correction *(p* < 0.05). **c**, **d**, Time course of germination rate (**c**) and sporulation rate (**d**) in media with and without 100 nM GR24. Dashed, solid and dotted lines indicate trials 1 to 3, respectively. Asterisks above graphs indicate significant differences among treatments in each trial using Fisher’s exact test with Bonferroni correction. ***, *p* < 0.001; *, 0.01 ≤*p* < 0.05; ns, not significant. DAI, days after incubation. **e**, Numbers of secondary spores in cultures with different concentrations of methyl jasmonate (MeJA) and GR24 at 6 WAI. Diamonds indicate medians. *Different letters* indicate significant differences (Wilcoxon rank-sum test with Bonferroni correction, *p* < 0.05). **f**, Secondary spores generated from a single parent spore in T (500 μM Myr-K, 1 mg L^−1^ peptone), TG (500 μM Myr-K, 1 mg L^−1^ peptone and 100 nM GR24) and TGM media (500 μM Myr-K, 1 mg L^−1^ peptone, 100 nM GR24 and 1 μM MeJA) at 8 WAI. Arrowheads indicate parent spores. Large secondary spores were frequently observed in the medium containing both GR24 and MeJA (TGM) than in only GR24 (TG). Many brown spores were observed in TGM and TG media compared to the medium without the phytohormones (T). Bars = 500 μm. **g**, Size distribution of secondary spores produced from 8 parent spores in T, TG and TGM media at 8 WAI. **h**, Size distribution of daughter spores produced on the extraradical hyphae emerged from carrot hairy roots inoculated with a single spore. **i**, Percentage of large spores (≥70 μm in diameter) in secondary spores produced in T, TG and TGM media at 8 WAI. *Different letters* indicate significant differences among treatments in each trial using Fisher’s exact test with Bonferroni correction (*p* < 0.05). *p*-values are described in Supplementary Table 3.

### Methyl jasmonate strengthens the effects of GR24 and increases the number and size of secondary spores in asymbiotic culture

We tested the effect of MeJA on asymbiotic culture of *R. clarus*. In the absence of GR24 (T medium), MeJA showed no significant effect on sporulation rate and numbers of secondary spores (Fig. 2e and Supplementary Fig. 4). However, MeJA increased numbers of secondary spores in medium containing GR24 (TG medium). *R. clarus* finally produced medians of ~300 secondary spores with a maximum of over 700 at 6 WAI in TG medium supplemented with 1 μM MeJA, hereinafter referred to as TGM medium. We also applied TGM medium to asymbiotic culture of the model AM fungus strain *R. irregularis* DAOM197198. The mean numbers of secondary spores per parent spore were 6–33 spores at 8 WAI (Supplementary Fig. 5), which was 4–22 fold compared with those in asymbiotic culture of *R. irregularis* reported previously^15^ but was inferior to *R. clarus* cultured in TGM medium.

### Time-lapse microscopy reveals development and structure details of *R. clarus* in asymbiotic culture

To reveal developmental patterns of *R. clarus* under asymbiotic conditions, we followed its growth and sporulation in TGM medium at two-hour intervals over time for 8 weeks by time-lapse microscopy (Supplementary Video 1). We summarised its developmental pattern in Supplementary Fig. 6. After the start of culture, parent spores produced germ tubes within one week. Germ tubes started to branch in four hours after germination in the fastest case and then produced RH. Tree-like branched hyphae were often observed from 1 to 2 days after germ tube emergence (DAG). Sometimes these branched hyphae continued to develop and formed large DPCs. DPC formation proceeded as follows. A single hypha, stemmed from RH laterally, generated short, thin hyphae at short intervals. Thereafter, these short hyphae elongated and branched many times (Supplementary Fig. 6). Large DPCs started appearing from one week after germ tube emergence (WAG) and were formed around parent spores and along elongating RH. The fastest secondary spores were formed apically or intercalarily along the lateral branches within 5 DAG. Development of secondary spores finished within 2 days after hyphal swelling (Supplementary Fig. 6). Once formed, these spores did not enlarge any more during the culture. The final diameter of the spores was positively correlated with the time required for the spore growth after hyphal swelling (Supplementary Fig. 6). *R. clarus* generated only small secondary spores in the early growth stage, thereafter the spore size was increased with days after germ tube emergence (Supplementary Fig. 6).

### The combination of GR24 and MeJA increases the proportion of large spores, but the spore size is still smaller than that of symbiotically-generated spores

We further analysed spore size in asymbiotic culture by comparing the size distributions between T, TG and TGM media (Fig. 2f–i). We applied machine-learning-based image analysis for enumerating secondary spores and measuring spore diameter. Spore size in any media had a right-skewed distribution with one peak around 40–50 μm in diameter (Fig. 2g). The size distribution in TGM medium had a higher right tail than that in the other media, indicating that the combination of GR24 and MeJA promoted the production of large spores. Large spores in TGM medium appeared brown in colour, similar to parent spores (Fig. 2f). Brown, large spores were also observed in TG medium, whereas the density was lower than that in TGM medium. In T medium, white, small spores were dominant. Spores larger than 70 μm in diameter seemed brown colour. The proportion of this size class was highest in TGM medium (Fig. 2i).

Next, we compared asymbiotically-generated secondary spores with daughter spores that were symbiotically produced by *in vitro* monoxenic culture. In monoxenic culture, a single spore was inoculated to carrot hairy roots on the M medium. During the 8-week cultivation period, daughter spores were produced on the extraradical hyphae emerged from the roots. Numbers of daughter spores in the monoxenic culture were almost the same as those of secondary spores formed in TGM medium (Supplementary Fig. 7). Size distribution of the daughter spores displayed two peaks (Fig. 2h). The lower peak around 50 μm was corresponded to the mode length of spore diameter in asymbiotic cultures, while the higher peak around 150 μm was not seen in any asymbiotic culture, indicating that secondary spores with an intermediate size between the two peaks were increased by the application of GR24 and MeJA.

### Subculture of secondary spores produced by asymbiotic culture

To ascertain whether secondary spores of *R. clarus* produced by asymbiotic culture can be subcultured, a large single spore (> 100 μm in diameter) or a gel block containing 20–40 spores produced in TGM medium were transferred to a new medium and incubated for 6 weeks. Single-spore culture exhibited a reduced number of spores compared with the initial culture, while the sporulation rate was still very high (Supplementary Fig. 8). Spores produced in the subculture were white and smaller than that in the initial asymbiotic culture (Supplementary Fig. 8). In gel block culture, no significant differences were observed between its subculture and the initial asymbiotic culture in sporulation rate and numbers of secondary spores per parent spore (Supplementary Fig. 8). Brown spores of similar size formed in the initial asymbiotic culture were observed in the gel block subculture (Supplementary Fig. 8).

### Asymbiotically-generated spores can colonise plants and promote their growth

We investigated the ability of asymbiotically-generated spores (AS) of *R. clarus* to produce daughter spores after the successful colonisation in plant roots. A single AS produced in TGM medium or symbiotically-generated spore (SS) by *in vitro* monoxenic culture was inoculated to carrot hairy roots grown on the modified M medium.

Germination rate of AS and SS were about 90% or higher on the medium (Fig. 3a). The frequency of AS to produce daughter spores after emerging extraradical hyphae from roots tended to be lower than that of SS up to 4 WAI, but no significant difference was observed from 4–6 WAI. Finally, 70 to 90% of AS (82 to 95% of germinated AS) produced daughter spores at 8 WAI (Fig. 3b, c and Supplementary Fig. 9). This is a substantial improvement over the rate, less than 15%, in previous experiments^14,15^. Overall, the daughter spore production in monoxenic culture inoculated with AS was about 2 weeks behind in numbers of daughter spores compared to SS, although some AS produced only a few daughter spores (Fig. 3d). Numbers of daughter spores were significantly correlated with diameter of spores inoculated to carrot hairy roots (Fig. 3e).

**Figure 3.**
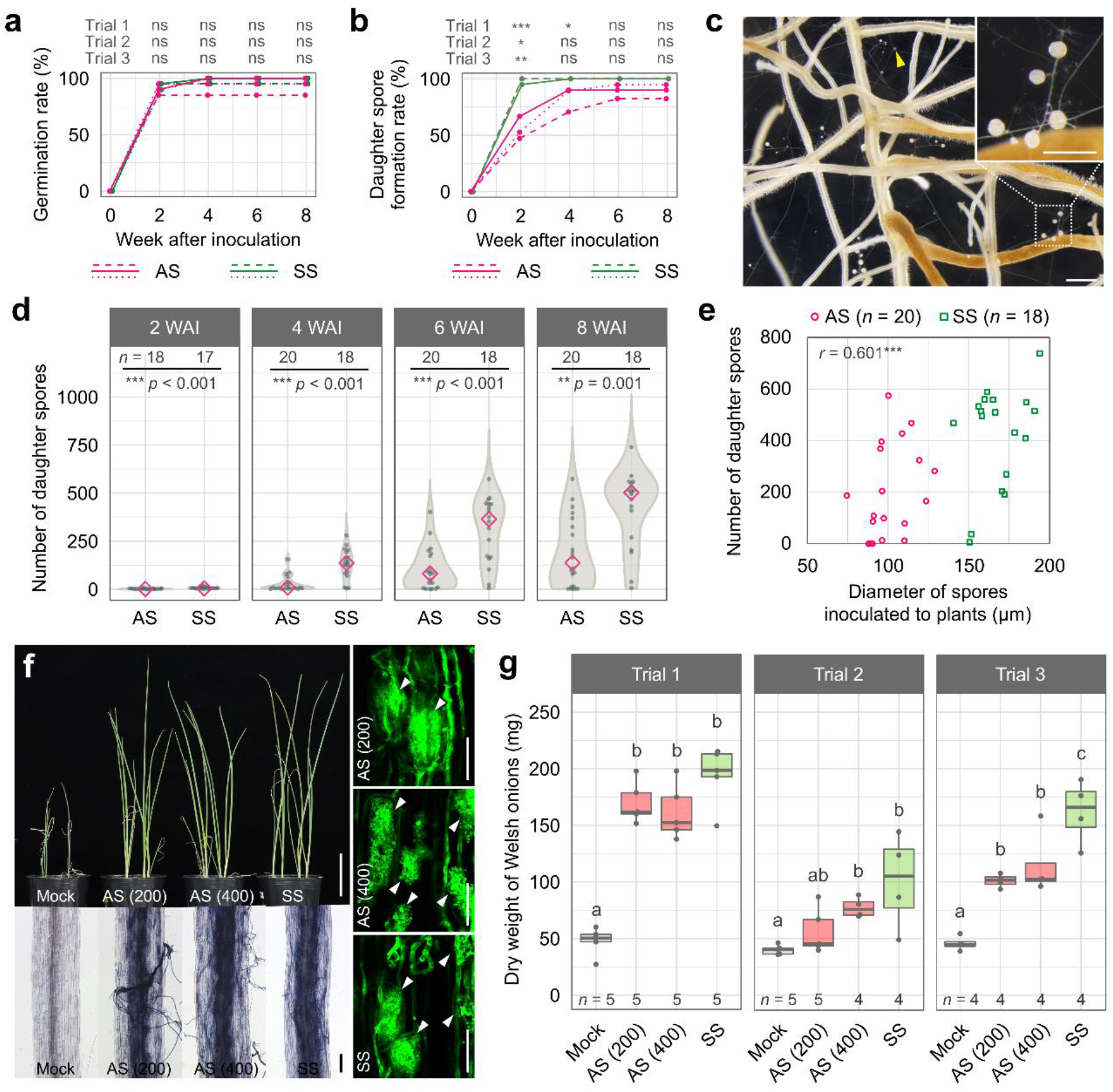
Inoculation of asymbiotically-generated spores to plants. **a–e**, Inoculation of asymbiotically-generated spores (AS) and symbiotically-generated spores (SS) to carrot hairy roots grown on modified M medium. AS and SS of *R. clarus* were prepared by asymbiotic culture in TGM medium and *in vitro* monogenic culture, respectively. Time-course of germination rate (**a**) and daughter spore formation rate calculated as percentage of spores that produce daughter spores after the successful colonisation relative to all spores inoculated to plants (**b**). Dashed, solid and dotted lines indicate trials 1 to 3, respectively. Asterisks above graphs indicate significant difference between treatments in each time point using Fisher’s exact test with Bonferroni correction. ***, *p* < 0.001; **, 0.001 ≤*p* < 0.01; *, 0.01 ≤*p* < 0.05; ns, not significant.*p*-values are described in Supplementary Table 3. **c**, Daughter spore formation in monoxenic culture, in which carrot hairy roots were inoculated with a single (arrowhead). Bars = 500 μm. **d**, Time course of daughter spore formation. Diamonds indicate medians. Asterisks above graphs indicate significant differences among treatments using Wilcoxon rank-sum test. **e**, Correlation between diameters of spores inoculated to carrot hairy roots and numbers of daughter spores at 8 weeks after inoculation (WAI). *r* value is Pearson’s correlation coefficient. ***, *p* < 0.001. **f–g**, Inoculation of AS and SS to Welsh onions in pots. **f**, Growth and root colonisation of Welsh onions inoculated with AM fungi at 8 WAI. Numbers in parentheses are numbers of spores for inoculation. Growth of Welsh onions (upper left, bar = 5 cm). AM fungal structures in roots stained with ink (bottom left, bar = 200 μm) and wheat germ agglutinin conjugated with Oregon green 488 (right, bars = 50 μm). Arrowheads show arbuscules. **g**, Dry weights of Welsh onion shoot at 8 WAI. Data were transformed as log_10_ (*x*). The upper and lower side of boxes show the 25 and 75% quantiles, bars inside boxes indicate medians. Whiskers indicate 1.5 times the interquartile range. *Different letters* indicate significant differences among treatments in each trial using Tukey’s honest significant difference test (*p* < 0.05).

We also inoculated AS and SS to Welsh onions in pots. AM fungal colonisation and arbuscule formation were observed in both roots inoculated with AS and SS (Fig. 3f and Supplementary Fig. 9). The shoot dry weight of Welsh onions inoculated with SS and spore number 400 AS was significantly greater than non-inoculated plants, whereas spore number 200 AS inoculated to plants did not always affect the plant growth (Fig. 4g). AS inoculation did not achieve the same effect as SS inoculation, although a large variation was observed within the trials.

## Discussion

Jones (1924) reported AM fungi as obligate parasites based on the observation that these fungi eventually die when grown on agar, but grow well inside or closely attached to roots^26^. AM fungi are now recognised as obligate symbionts forming associations with most terrestrial plants^27^. Host-free culture of AM fungi more recently has been developed by supplying fatty acids, myristic acid and palmitoleic acid^14,15^. In the present study, we demonstrate that the combination of two phytohormones, strigolactone and MeJA, facilitates asymbiotical spore production of *R. clarus* in the presence of myristate and peptone. These asymbiotically-generated spores can be subcultured and promote plant growth after their inoculation, while the spore size was still smaller than that of spores produced by co-culture with plants.

As observed in *R. irregularis*^15^, myristate was essential for asymbiotic growth of *R. clarus* (Fig. 1a and Supplementary Fig. 1). GR24 drastically accelerated asymbiotic growth and sporulation of *R. clarus* in the presence of potassium myristate (Fig. 2a–d). Strigolactones are known to be plant-derived signal molecules that activate mitochondrial activity of AM fungi and to promote hyphal elongation and branching during the pre-symbiotic phase^19–21^. The rapid colony expansion stimulated by GR24 in TG or TGM medium may further augment the amount of nutrients absorbed into its hyphae, leading to the early spore formation and mass production of secondary spores. GR24 and MeJA acted synergistically on the increase in numbers and size of secondary spores (Fig. 2e–i). Plant genes related to jasmonic acid biosynthesis are found to be specifically expressed in arbuscule-containing cortical cells of roots^23,24^. However, the crosstalk between jasmonic acid and strigolactone signalling in AM symbiosis and the direct effect of jasmonic acid on AM fungal growth are unknown. Future study is needed to elucidate roles of jasmonic acid in asymbiotic growth of AM fungi.

Time-lapse image analysis revealed development pattern of asymbiotic growth of *R. clarus* (Supplementary Video 1). In its early growth stage, small DPC structures like branched absorbing structure (BAS)^28^, which is observed in extraradical hyphal networks in monoxenic culture, were generated along RH. Thereafter, large DPCs with longer and more branched hyphae were formed. DPC was first described in co-culture of *R. irregularis* with *P. validus*^12^, and also reported in asymbiotic culture supplemented with myristate^15^, indicating that DPC is a specific fungal structure in *Rhizophagus* spp. cultured without host plants. In *R. irregularis* cultured in TGM medium, a relatively large DPC was formed in the vicinity of a parent spore, and then small DPCs, identified as BAS-like structures in the previous report^15^, developed along RH. In contrast, large DPCs of *R. clarus* were developed not only around parent spores but also away from the spores (Supplementary Video 1). Fatty acids are absorbed from branched hyphae like DPC, and part of the fatty acids or lipids are transferred to newly formed secondary spores via RH in *R. irregularis*^15^. *R. clarus* can produce many large spores in TGM medium possibly owing to massive uptake of myristate via many large DPC formed at its later growth stage. The final size of asymbiotically-generated spores was smaller than that of spores produced by co-culture with plants. The spore development was finished in a very shorter period, within 2 days after hyphal swelling (Supplementary Fig. 6). In monoxenic culture of *R. irregularis*, it takes 30 to 60 days to reach mature spore size^29^. Maturation of spores in asymbiotic cultures might require other plant or environmental factors.

For developing axenic culture system of AM fungi, subculture and inoculum potential of fungal materials are important aspects. *R. irregularis* spores generated in palmitoleic acid-containing medium without hosts can germinate and form new secondary spores in the second culture, but the spore productivity was low^14^. In TGM medium, *R. clarus* can be subcultured from a single spore and multiple spores in a gel block. In particular, the agar block subculture showed almost the same level of spore production as the initial culture. This finding prompts us in the future to test asymbiotically continuous culture of *R. clarus*. Secondary spores produced in TGM medium had the ability to colonise host plants. Although inoculation of AS promoted the growth of Welsh onions, the mycorrhizal effect was lower than SS. The lower effect of AS may be related to the small spore size because inoculation of AS to carrot hairy roots showed the delayed establishment of AM symbiosis.

### Perspective

This study raises three interesting perspectives for this awkward microorganism. The first is elucidation of differences between fungal species. *R. irregularis* forms more spores on TGM medium (Supplementary Fig. 5) than medium without hormones which is previously reported^15^, but produced secondary spores are fewer than in *R. clarus*. Asymbiotic growth of *R. irregularis* might also be stimulated by GR24 similar to *R. clarus*, but with less impact. It implies that pathways involved in AM fungal growth could be different even within the same genus, *Rhizophagus*. The genomes of both *R. irregularis* and *R. clarus* have been sequenced, allowing investigation of lineage-specific missing genes that might suggest suitable culture methods for individual fungi by comparative genome analysis. The second aspect is the contribution to biochemical analysis. Collecting large amounts of chemical constituents of AM fungi is difficult. Easy, large-scale culture of *R. clarus* will encourage investigation and biochemical analysis of fungal effectors, transporters, small signalling molecules such as Myc-LCO, as well as identification of strigolactone and MeJA binding proteins. The third contribution involves field research. AS of *R. clarus* significantly promotes Welsh onion growth (Fig. 3f, g). Realising a large-scale culture system at low cost will allow field inoculation experiments without the use of plant roots.

Ancestors of AM fungi were likely autonomous microorganisms. AM fungi lost autonomy in the process of evolution over 400 million years of coexistence with host plants^30^and became obligate symbionts. There is no way to know the appearance and behaviour of the ancient AM free-living fungi. However, the movie (unique growth and proliferation of *R. clarus*, Supplementary Video 1) shown in this study may reflect, to some extent, the appearance of AM fungi hundreds of millions of years ago.

## Materials and Methods

### Fungal materials

The fungal strain *R. clarus* HR1 was originally isolated from the quarry in Hazu, Aichi, Japan^31^ and is available from the NARO Genebank (https://www.gene.affrc.go.jp/index_en.php) as *R. clarus* MAFF520076. Sterile spores were collected from *in vitro* monoxenic culture of *R. clarus* in association with carrot hairy roots^20^ grown on M medium^32^, solidified with 0.4% of gellan gum (FUJIFILM Wako) instead of agar, at 28 °C in the dark. *R. irregularis* DAOM197198 (or DAOM181602, another voucher number for the same fungus) was purchased from Premier Tech, Canada.

### Preparation of media for asymbiotic culture

Unless otherwise noted, all chemicals were purchased from FUJIFILM Wako. Stock solutions of 100 mM potassium myristate, 100 mM potassium palmitate and 10 mM 2OH-TDA (Cayman Chemical) were prepared by dissolving in water and sterilised by filtration. Potassium palmitoleate was prepared by adding 10 mM potassium hydroxide to palmitoleic acid (Sigma-Aldrich), which was finally adjusted to 10 mM by water. Bacto™ Peptone (BD), Bacto™ Yeast Extract (BD), Bacto™ Malt Extract (BD) were dissolved in water and autoclaved at 121 °C for 20 min. Synthetic Complete (KAISER) DROP-OUT: COMPLETE (SC DROP-OUT, FORMEDIUM) was dissolved in water and sterilised by filtration. Strigolactone GR24 was synthesised as previously described^33^ and dissolved in acetone for a 0.1 mM stock solution. Methyl jasmonate was dissolved in ethanol for a 1 mM stock solution. For preparation of culture media, all compounds except vitamins listed in Supplementary Table 1 were added in water and sterilised by autoclaving. Vitamins and supplements of fatty acid salts, organic nitrogen sources and phytohormones were added to the autoclaved solution under sterile conditions. A stock solution of fatty acid salt was slowly added with stirring. Media were poured into 90-mm Petri dishes and solidified by cooling. T medium was a base medium (Supplementary Table 1) containing 500 μM potassium myristate and 1.0 mg L^−1^ peptone. TG and TGM media were prepared by adding 100 nM GR24 and both 100 nM GR24 and 1 μM MeJA to T medium, respectively.

### Asymbiotic culture

Parent spores of *R. clarus* as seed fungus for asymbiotic culture were collected from *in vitro* monoxenic culture, cultivated for at least three months after inoculation (Supplementary Table 2). Spores were separated from extraradical hyphae with forceps and scalpels under an SZX7 stereomicroscope in a laminar flow cabinet. Collected spores were pooled in sterile water, then placed separately on a solid medium using a pipette with water droplets to prevent dryness under sterile conditions. Large (120-220 μm *R. clarus* and 80-140 μm diameter *R. irregularis*) and coloured spores were chosen. Four to ten parent spores of *R. clarus* or *R. irregularis* were placed on a solid medium in Petri dish (numbers of parent spores are described in Supplementary Table 2). AM fungi were grown at 28 °C in the dark for 6 or 8 weeks. Numbers of secondary spores and spore diameters were measured under an SZX7 stereomicroscope (Olympus).

### Time-lapse microscopy

Three or five parent spores of *R. clarus* were placed in the centre of a TGM plate (containing 400 or 500 μM Myr-K). The time-lapse images were captured every 2 hours at 28 °C in the dark for 2 months using an SZX7 stereomicroscope equipped with a DP21 digital camera system (Olympus). All images were acquired under constant intensity transmitted light. Time-lapse images were generated by integrating individual digital images with ImageJ^34^ Spore diameters were measured using ImageJ.

### Machine-learning-based image analysis of asymbiotically-generated spores

Machine-learning-based image analysis using Ilastik software (https://www.ilastik.org/index.html) was applied for enumerating secondary spores in asymbiotic culture at 8 WAI. Solid medium containing fungal materials was cut using forceps and scalpels, and then incubated in a three times volume of citrate buffer (1.7 mM citric acid and 8.3 mM trisodium citrate, pH 6.0) for over 2 hours at room temperature. After centrifuge at 3,200 ×*g* for 15 min at room temperature, fungal pellets were suspended in 500 μl water and transferred to a new tube. Fungal materials were shredded with an ultrasonic processor (SONICS Vibra-Cell VCX130) as follows: 40% amplitude, 3 sec ON/4 sec OFF pulses for 1-5 min, 2 mm diameter probe. After centrifuge, 300 μl of the supernatant was removed, and fungal pellet was suspended. An aliquot containing ~300 spores was placed on a Petri dish. Spores were dispersed using forceps and allowed to settle down for over one hour. Digital images of spores were captured using an SZX7 stereomicroscope with a digital camera system (DP73, Olympus) under transmitted light. Spore counting by machine-learning was performed using the pixel classification workflow of Ilastik. First, training of a classifier that can separate spores from background was done. Next, individual spore images were extracted by image binarization using simple segmentation function. Numbers of spores were counted using the binarised images by ImageJ^34^. Spore diameter was estimated from maximum feret diameter measured by ImageJ. Spores smaller than 30 μm in diameter were excluded from the analysis.

### Subculture of asymbiotically-generated spores

*R. clarus* secondary spores with diameter of > 100 μm generated by asymbiotic culture of at least 3 months were used for subcultures (Supplementary Table 2). A single spore or an agar block (5 × 5 mm square) containing 20–40 secondary spores was transferred to a new TGM medium using forceps and scalpels. AM fungi were incubated at 28 °C in the dark for 6 weeks. Numbers of newly formed secondary spores were counted under an SZX7 stereomicroscope.

### Inoculation test of asymbiotically-generated spores to plants

For inoculation test of *R. clarus* to carrot hairy roots, spores produced by asymbiotic and *in vitro* monoxenic culture for at least 3 months were used as inoculum (Supplementary Table 2). *R. clarus* secondary spores with diameter of > 70 μm generated by asymbiotic culture were chosen. A single spore was harvested using forceps and scalpels under an SZX7 stereomicroscope and placed on the vicinity of carrot hairy roots grown on M medium. The production of daughter spores on extraradical hyphae emerging from hairy roots was observed under the stereomicroscope.

For inoculation to Welsh onion, *R. clarus* spores produced by asymbiotic and *in vitro* monoxenic culture for at least 4 months were used as inoculum. Secondary spores formed in asymbiotic culture were collected after melting solid medium by citrate buffer as described above. Seeds of Welsh onion cultivar Asagi-kujo-hosonegi (TOHOKU SEED, Japan) were sown in sterile soil (1:1 mixture of Akadamatsuchi and Kanumatutipumice soil) and incubated in the dark for 5–6 days at 24 °C. One day before inoculation, seedlings were transferred to long-day conditions [16 hours of light and 8 hours of dark]. Spores were mixed in 160 ml of humic soil (Kurotsuchi) that was supplemented with 50 ml of modified Long Ashton liquid medium^35^ containing 20 μM phosphate (Supplementary Table 4) after autoclaved at 121°C for 40 min. Four seedlings were transplanted in each pot (7 cm in diameter) filled with the inoculated soil and cultivated at 24 °C for 8 weeks. After harvesting, plant shoots were dried at 80 °C for four days or 50 °C for 2 weeks and weighed with an analytical balance.

### Observation of AM fungal structures in roots

AM roots were stained using ink (Fountain Pen ink 4001 brilliant black, Pelikan) and wheat germ agglutinin conjugated with Oregon Green 488 (Thermo Fisher Scientific) as described by Takeda et al.^36^. Images of ink and wheat germ agglutinin staining were acquired using a BX50 microscope equipped with a DP73 digital camera system (Olympus) and a Nikon A1 confocal laser scanning microscope, respectively.

### Statistical analysis

All statistical analyses were performed using R software (version 4.0.2). To examine the differences among experimental groups, data were analysed with Wilcoxon rank-sum test and Fisher’s exact test, as appropriate. For multiple comparisons, *p*-values were corrected by the Bonferroni method.

## Acknowledgements

We thank Prof. Masanori Saito for kindly advise. We also thank Yumi Yoshinori, Yuuko Ogawa and Asami Tokairin for experimental supports. The modified low-P Long Ashton medium was developed by Ei-ichi Murakami. This research was supported by the Model Plant Research Facility of National Institute for Basic Biology for lending equipment (NIBB). Confocal images were acquired at Spectrography and Bioimaging Facility, NIBB Core Research Facilities. This work was supported by ACCEL (JPMJAC1403) form the Japan Science and Technology Agency (to T.E., K.S., K.A. and M.K.).

## Author Contributions

Asymbiotic culture and *in vitro* monoxenic culture of AM fungi were performed by S.T. and K.H. A survey of organic nitrogen sources was done by H.K. Analyses of asymbiotically-generated spores were performed by K.H. Time-lapse images were acquired by Y.K. and K.H. and analysed by Y.K. Inoculation tests of AM fungal spores into Welsh onions were performed by S.T., K.H. and K.Y. Data was statistically analysed by T.M. The strigolactone GR24 was synthesised by K.A. K.H., K.S., Y.K. and M.K. wrote the manuscript. T.E., K.S., K.A. and M.K. planed this research. M.K. supervised this study.

## Competing Financial Interests Statement

The authors declare no competing financial interests.

## Notes

### Competing Interest Statement

The authors have declared no competing interest.

## References

1. Sally Smith & David Read. Mycorrhizal Symbiosis - 3rd Edition. (Elsevier, 2008).

2. Bonfante, P. & Genre, A. Plants and arbuscular mycorrhizal fungi: an evolutionary-developmental perspective. Trends Plant Sci. 13, 492–498 (2008).

3. Bravo, A., Brands, M., Wewer, V., Dörmann, P. & Harrison, M. J. Arbuscular mycorrhiza-specific enzymes FatM and RAM2 fine-tune lipid biosynthesis to promote development of arbuscular mycorrhiza. New Phytol. 214, 1631–1645 (2017).

4. Keymer, A. et al. Lipid transfer from plants to arbuscular mycorrhiza fungi. eLife 6,.

5. Luginbuehl, L. H. et al. Fatty acids in arbuscular mycorrhizal fungi are synthesized by the host plant. Science 356, 1175–1178 (2017).

6. Jiang, Y. et al. Plants transfer lipids to sustain colonization by mutualistic mycorrhizal and parasitic fungi. Science 356, 1172–1175 (2017).

7. Kobayashi, Y. et al. The genome of *Rhizophagus clarus* HR1 reveals a common genetic basis for auxotrophy among arbuscular mycorrhizal fungi. BMC Genomics 19, 1–11 (2018).

8. Tisserant, E. et al. Genome of an arbuscular mycorrhizal fungus provides insight into the oldest plant symbiosis. Proc. Natl. Acad. Sci. U. S. A. 110, 20117–20122 (2013).

9. Wewer, V., Brands, M. & Dörmann, P. Fatty acid synthesis and lipid metabolism in the obligate biotrophic fungus *Rhizophagus irregularis* during mycorrhization of *Lotus japonicus*. Plant J. 79, 398–412 (2014).

10. Sun, X. et al. Genome and evolution of the arbuscular mycorrhizal fungus *Diversispora epigaea* (formerly *Glomus versiforme)* and its bacterial endosymbionts. New Phytol. 221, 1556–1573 (2019).

11. Tang, N. et al. A survey of the gene repertoire of *Gigaspora rosea* unravels conserved features among *Glomeromycota* for obligate biotrophy. Front. Microbiol. 7, (2016).

12. Hildebrandt, U., Janetta, K. & Bothe, H. Towards growth of arbuscular mycorrhizal fungi independent of a plant host. Appl. Environ. Microbiol. 68, 1919–1924 (2002).

13. Hildebrandt, U., Ouziad, F., Marner, F.-J. & Bothe, H. The bacterium *Paenibacillus validus* stimulates growth of the arbuscular mycorrhizal fungus *Glomus intraradices* up to the formation of fertile spores. FEMS Microbiol. Lett. 254, 258–267 (2006).

14. Kameoka, H. et al. Stimulation of asymbiotic sporulation in arbuscular mycorrhizal fungi by fatty acids. Nat. Microbiol. 4, 1654–1660 (2019).

15. Sugiura, Y. et al. Myristate can be used as a carbon and energy source for the asymbiotic growth of arbuscular mycorrhizal fungi. Proc. Natl. Acad. Sci. (2020).

16. Gianinazzi-Pearson, V., Séjalon-Delmas, N., Genre, A., Jeandroz, S. & Bonfante, P. Plants and arbuscular mycorrhizal fungi: Cues and communication in the early steps of symbiotic interactions. Adv. Bot. Res. 46, 181–219 (2007).

17. Nagata, M. et al. Enhanced hyphal growth of arbuscular mycorrhizae by root exudates derived from high R/FR treated *Lotus japonicus*. Plant Signal. Behav. 11, e1187356 (2016).

18. Bücking, H. et al. Root exudates stimulate the uptake and metabolism of organic carbon in germinating spores of *Glomus intraradices*. New Phytol. 180, 684–695 (2008).

19. Akiyama, K., Matsuzaki, K. & Hayashi, H. Plant sesquiterpenes induce hyphal branching in arbuscular mycorrhizal fungi. Nature 435, 824–827 (2005).

20. Tsuzuki, S., Handa, Y., Takeda, N. & Kawaguchi, M. Strigolactone-induced putative secreted protein 1 is required for the establishment of symbiosis by the arbuscular mycorrhizal fungus *Rhizophagus irregularis*. Mol. Plant-Microbe Interact. 29, 277–286 (2016).

21. Besserer, A. et al. Strigolactones stimulate arbuscular mycorrhizal fungi by activating mitochondria. PLoS Biol. 4, (2006).

22. Wu, Q.-S. et al. Mycorrhiza alters the profile of root hairs in trifoliate orange. Mycorrhiza 26, 237–247 (2016).

23. Hause, B., Maier, W., Miersch, O., Kramell, R. & Strack, D. Induction of jasmonate biosynthesis in arbuscular mycorrhizal barley roots. Plant Physiol. 130, 1213–1220 (2002).

24. Hause, B. & Schaarschmidt, S. The role of jasmonates in mutualistic symbioses between plants and soil-born microorganisms. Phytochemistry 70, 1589–1599 (2009).

25. Nagahashi, G. & Douds, D. D. The effects of hydroxy fatty acids on the hyphal branching of germinated spores of AM fungi. Fungal Biol. 115, 351–358 (2011).

26. Jones, F. R. A mycorrhizal fungus in the roots of legumes and some other plants. J. Agric. Res. 29, 459–470 (1924).

27. Brundrett, M. C. & Tedersoo, L. Evolutionary history of mycorrhizal symbioses and global host plant diversity. New Phytol. 220, 1108–1115 (2018).

28. Bago, B., Azcon-Aguilar, C., Goulet, A. & Piche, Y. Branched absorbing structures (BAS): a feature of the extraradical mycelium of symbiotic arbuscular mycorrhizal fungi. New Phytol. 139, 375–388 (1998).

29. Marleau, J., Dalpé, Y., St-Arnaud, M. & Hijri, M. Spore development and nuclear inheritance in arbuscular mycorrhizal fungi. BMC Evol. Biol. 11, 1–11 (2011).

30. Taylor, T. N., Remy, W., Hass, H. & Kerp, H. Fossil arbuscular mycorrhizae from the Early Devonian. Mycologia 87, 560–573 (1995).

31. Maki, T., Nomachi, M., Yoshida, S. & Ezawa, T. Plant symbiotic microorganisms in acid sulfate soil: significance in the growth of pioneer plants. Plant Soil 310, 55–65 (2008)

32. Bécard, G. & Fortin, J. A. Early events of vesicular–arbuscular mycorrhiza formation on Ri T-DNA transformed roots. New Phytol. 108, 211–218 (1988).

33. Mangnus, E. M., Dommerholt F. Jan., De Jong, R. L. P. & Zwanenburg, Binne. Improved synthesis of strigol analog GR24 and evaluation of the biological activity of its diastereomers. J. Agric. Food Chem. 40, 1230–1235 (1992).

34. Schneider, C. A., Rasband, W. S. & Eliceiri, K. W. NIH Image to ImageJ: 25 years of Image Analysis. Nat. Methods 9, 671–675 (2012).

35. Hewitt, E. J. Sand and water culture methods used in the study of plant nutrition. Commonwealth Agriculture Bureaux. (1966).

36. Takeda, N. et al. Gibberellins interfere with symbiosis signaling and gene expression and alter colonization by arbuscular mycorrhizal fungi in *Lotus japonicus*. Plant Physiol. 167, 545–557 (2015).

