## Supplementary Figures and Tables. for "Asymbiotic mass production of the arbuscular mycorrhizal fungus *Rhizophagus clarus*"

**Title**

Supplementary Figures 1–9

Supplementary Tables 1–4

Supplementary Video 1

A video of *R. clarus* HR1 developmental pattern, which generated by images taken at two-hour intervals over time for 8 weeks by time-lapse microscopy. The number at the bottom left indicates days after germ tube emergence.

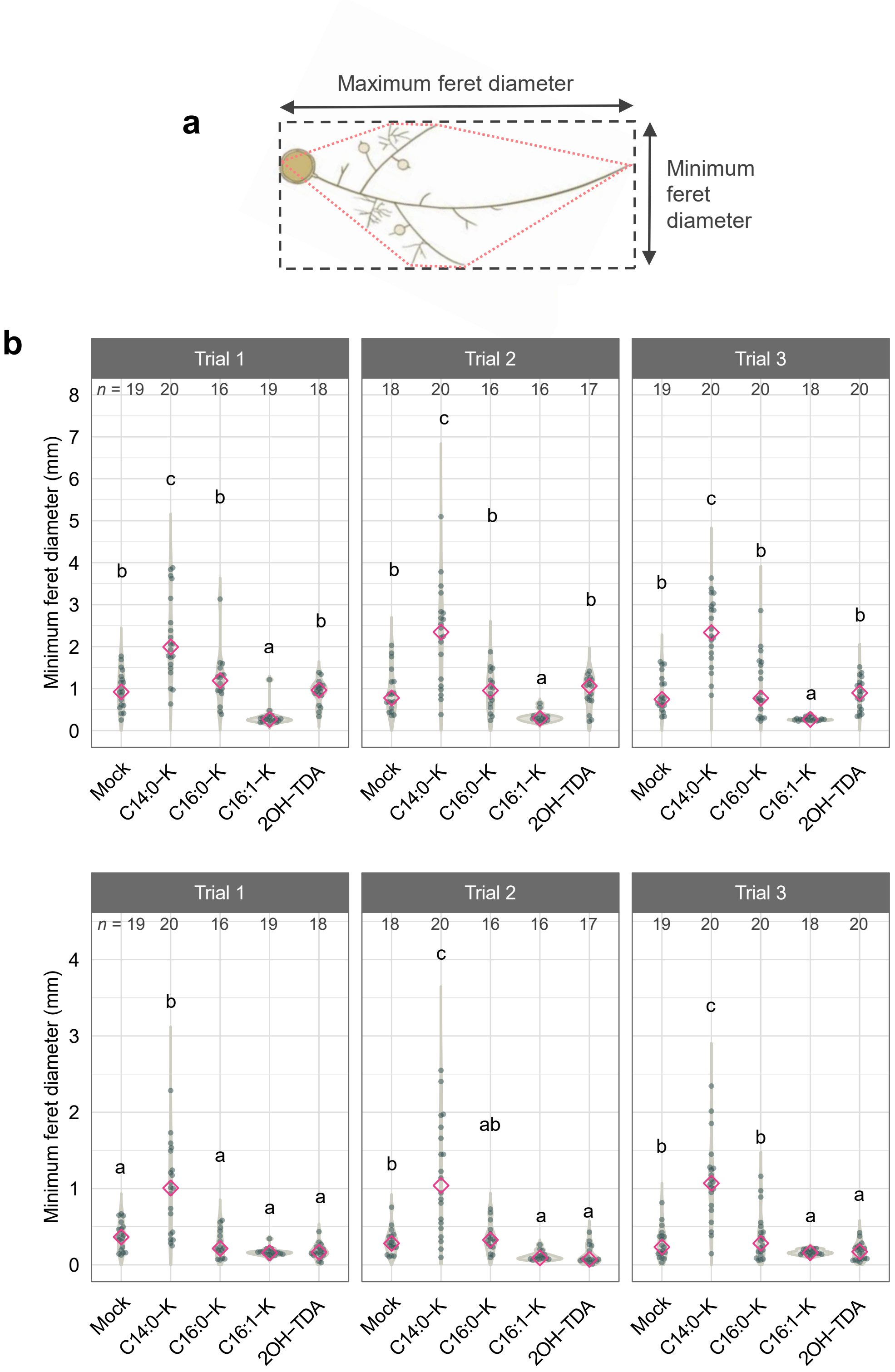


**Supplementary Figure 1. Effects of fatty acids on asymbiotic culture for *R. clarus*.** **a**, The outline of the measuring method using maximum and minimum feret diameters (explained in dashed box) for fungal spread hyphae. Feret diameters were analysed using Image J software from a polygon drawn by connecting the tips of hyphae with straight lines (shown in dotted red line). **b**, Maximum and minimum feret diameter of *R. clarus* in conditions supplemented with 100 µM fatty acid at 6 weeks after incubation (WAI). Diamonds indicate medians. Statistical significance was calculated using the Wilcoxon rank-sum test with Bonferroni correction. *Different letters* indicate significant differences ($p < 0.05$). $p$-values are described in Supplementary Table 3. C14:0-K, potassium myristate. C16:0-K, potassium palmitate. C16:1-K, potassium palmitoleate. 2OH-TDA, 2-hydroxytetradecanoic acid.

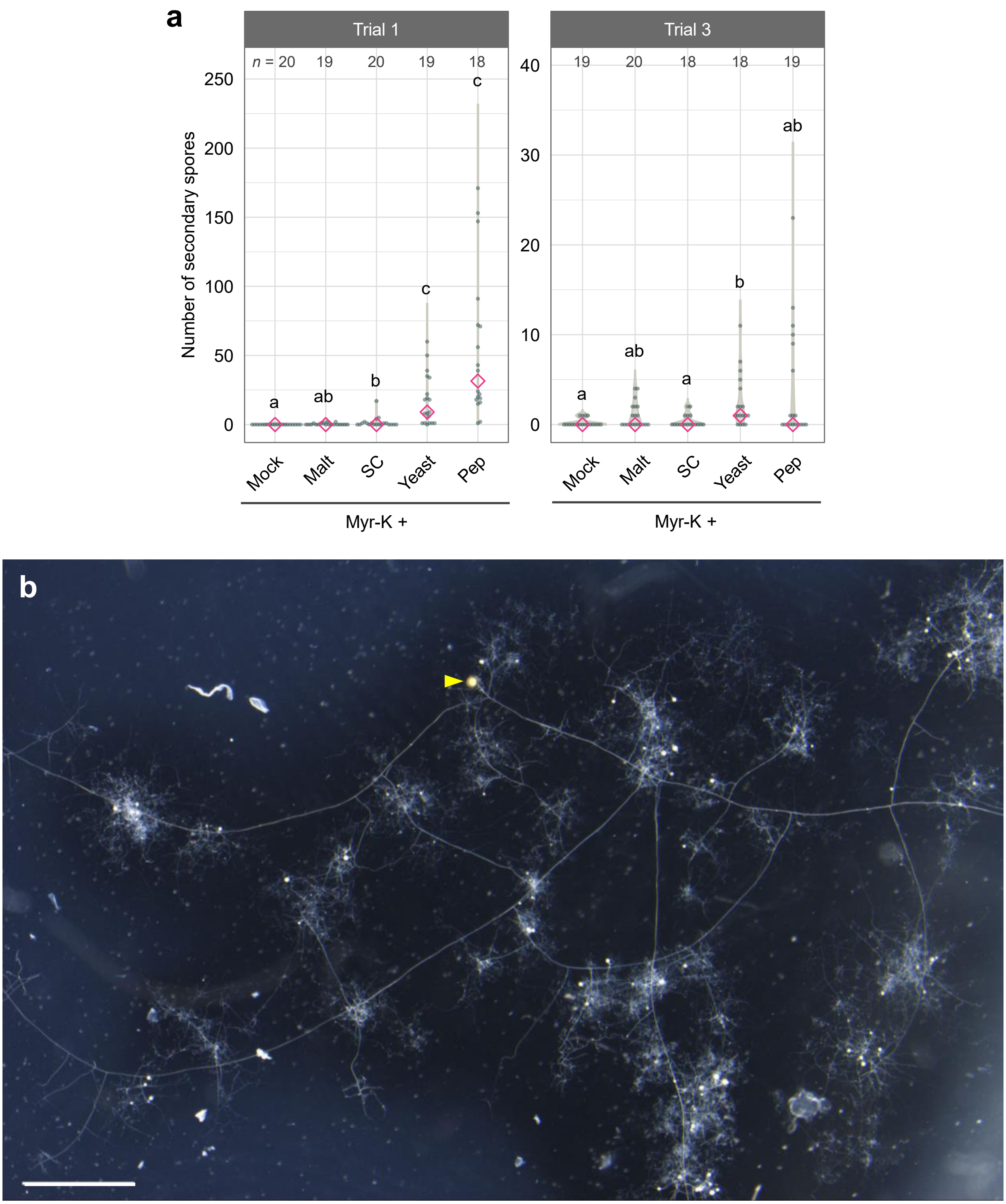


**Supplementary Figure 2. Effects of organic nitrogen on asymbiotic culture for *R. clarus*. a**, Another two trials of Fig. 1d (trial 2); numbers of secondary spores in medium supplemented with 0.2 mg $L^{-1}$ organic nitrogen in the presence of Myr-K at 6 WAI. Diamonds indicate medians. Statistical significance was calculated using the Wilcoxon rank-sum test with Bonferroni correction. *Different letters* indicate significant differences ($p < 0.05$). $p$-values are described in Supplementary Table 3. Malt, malt extract. Pep, peptone. SC, SC dropout. Yeast, yeast extract. **b**, Low magnification image of *R. clarus* cultured in the medium containing Myr-K and peptone in Fig. 1a. An arrowhead, a parent spore. Bar = 2 mm.

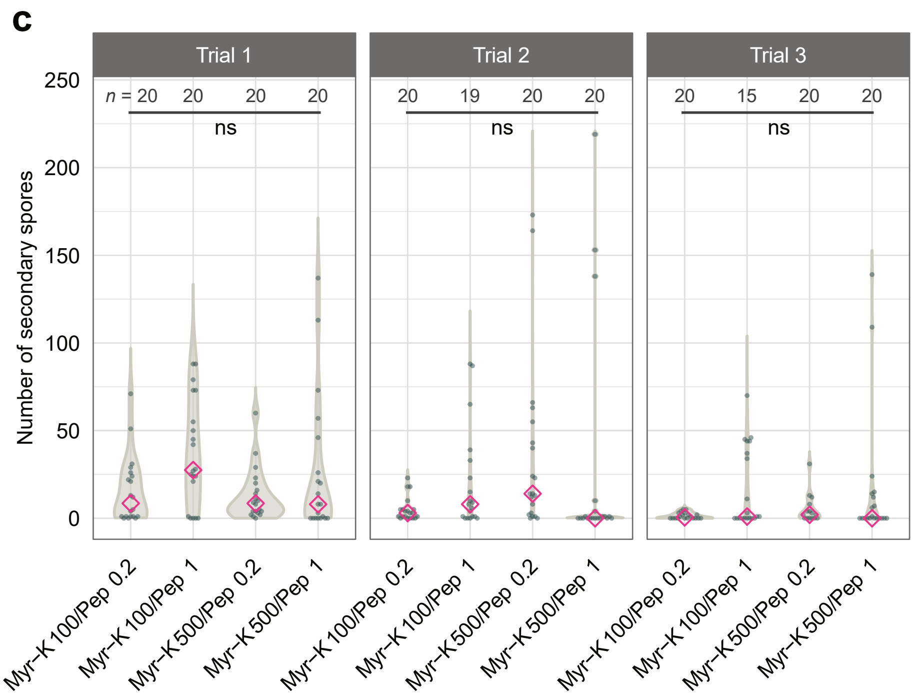


**c**, Comparison of secondary spore number among different combinations of Myr-K (100 or 500 µM) and peptone (1.0 or 0.2 mg $L^{-1}$) at 6 WAI. Diamonds indicate medians. Statistical significance was calculated using the Wilcoxon rank-sum test with Bonferroni correction ($p < 0.05$). ns, not significant. *p*-values are described in Supplementary Table 3.

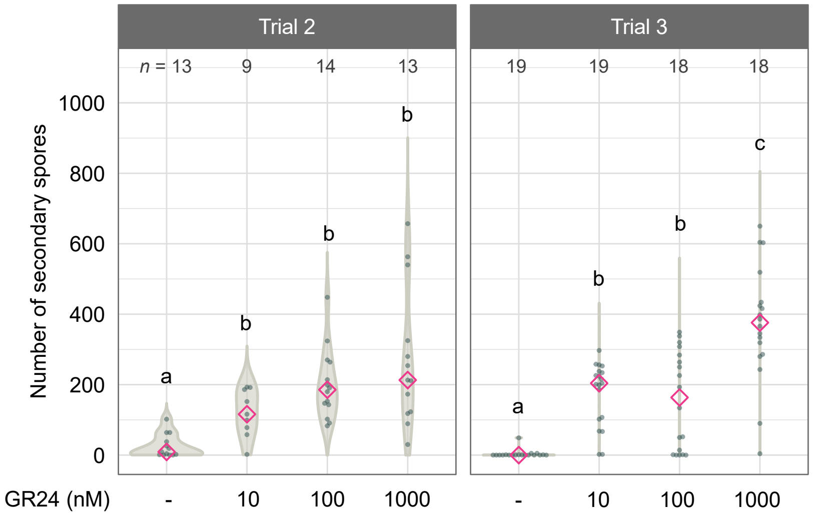


**Supplementary Figure 3. Effects of GR24 on asymbiotic culture for *R. clarus*.** Another two trials of Fig. 2b (trial 1); numbers of secondary spores in cultures with different concentration of GR24 in the presence of 500 µM Myr-K and 1 mg $L^{-1}$ peptone at 6 WAI. Diamonds indicate medians. Statistical significance was calculated using the Wilcoxon rank-sum test with Bonferroni correction. *Different letters* indicate significant differences ($p < 0.05$). *p*-values are described in Supplementary Table 3.

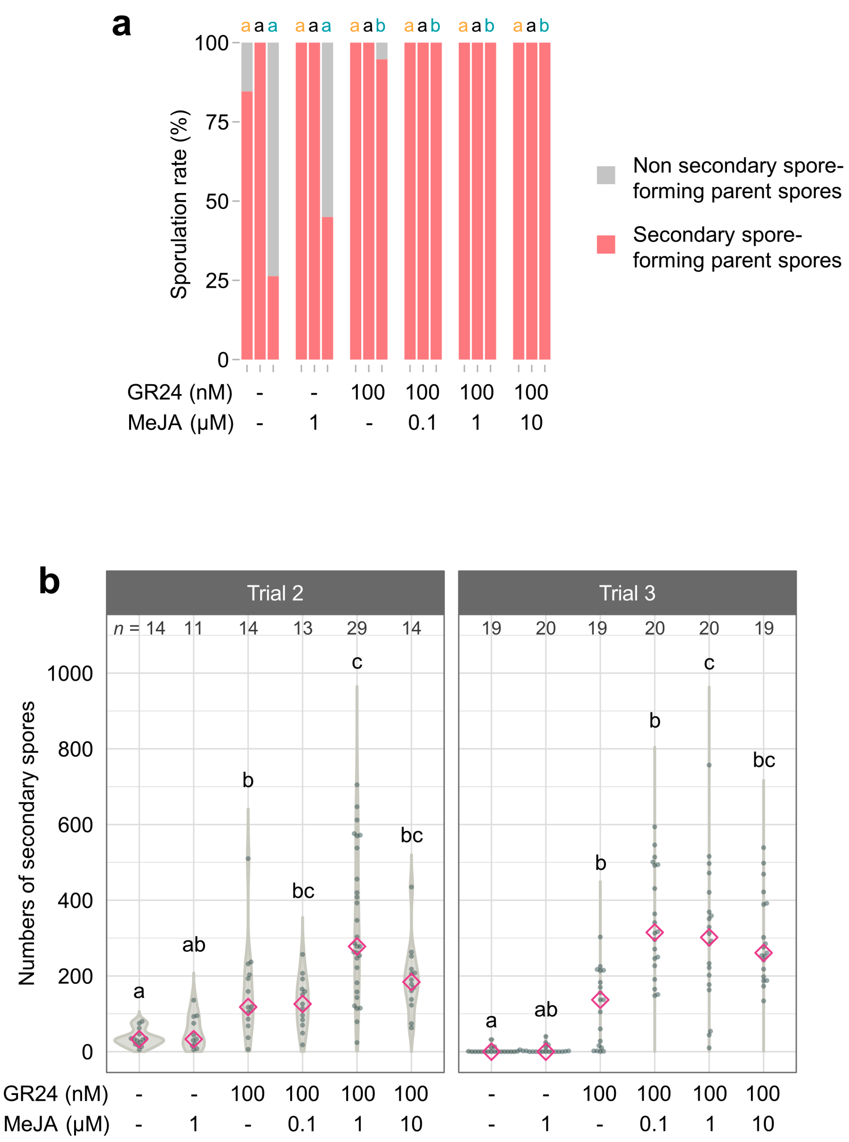


**Supplementary Figure 4. Effects of GR24 and methyl jasmonate on asymbiotic culture for *R. clarus*.** **a**, Sporulation rates, percentage of secondary spore-forming parent spores relative to germinated patent spores, in medium containing GR24 or MeJA or both at 6 WAI. Three bars in each condition are trial 1 to 3, respectively from the left. *Different letters* above of the graph indicate significant differences among treatments in each trial using Fisher's exact test with Bonferroni correction ($p < 0.05$). **b**, Another two trials of Fig. 3a (trial 1); asymbiotic culture experiments with GR24 and MeJA as before. Diamonds indicate medians. Statistical significance was calculated using the Wilcoxon rank-sum test with Bonferroni correction. *Different letters* indicate significant differences ($p < 0.05$). *p*-values are described in Supplementary Table 3.

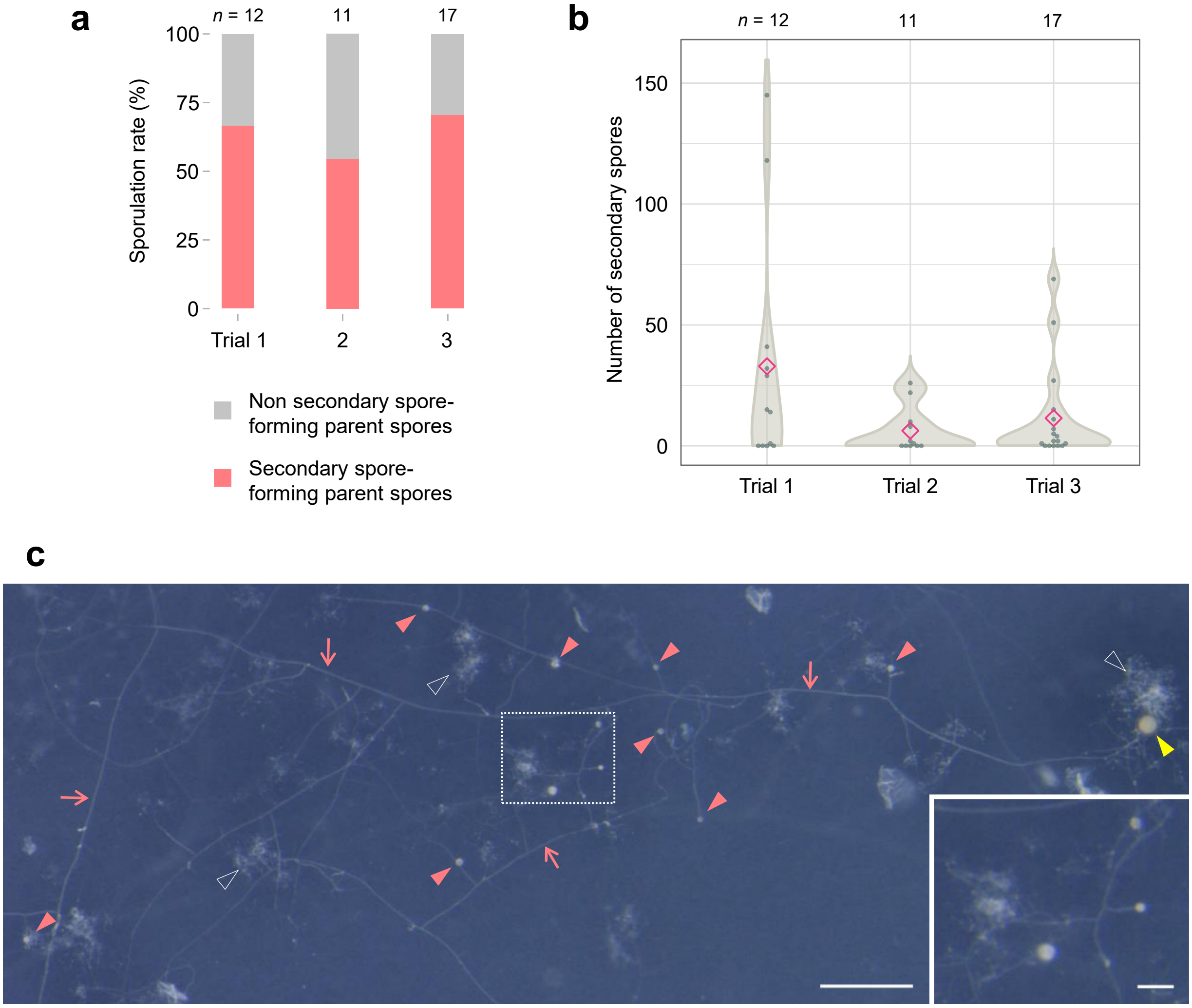


**Supplementary Figure 5. Asymbiotic culture of *R. irregularis* DAOM197198. a,** Sporulation rates , percentage of secondary spore-forming parent spores relative to germinated patent spores, in TGM medium at 8 WAI. **b**, Numbers of secondary spores produced in TGM medium at 8 WAI. Diamonds indicate means. **c**, *R. irregularis* secondary spores generated on TGM medium at 8 WAI. The inset is the magnified image of the dotted box. Yellow and pink arrowheads indicate a parent spore and newly generated secondary spores, respectively. Arrows indicate runner hyphae. Outlined arrowheads indicate small densely packed coils structures. Bars indicate 500 (large image) and 100 µm (inset), respectively.

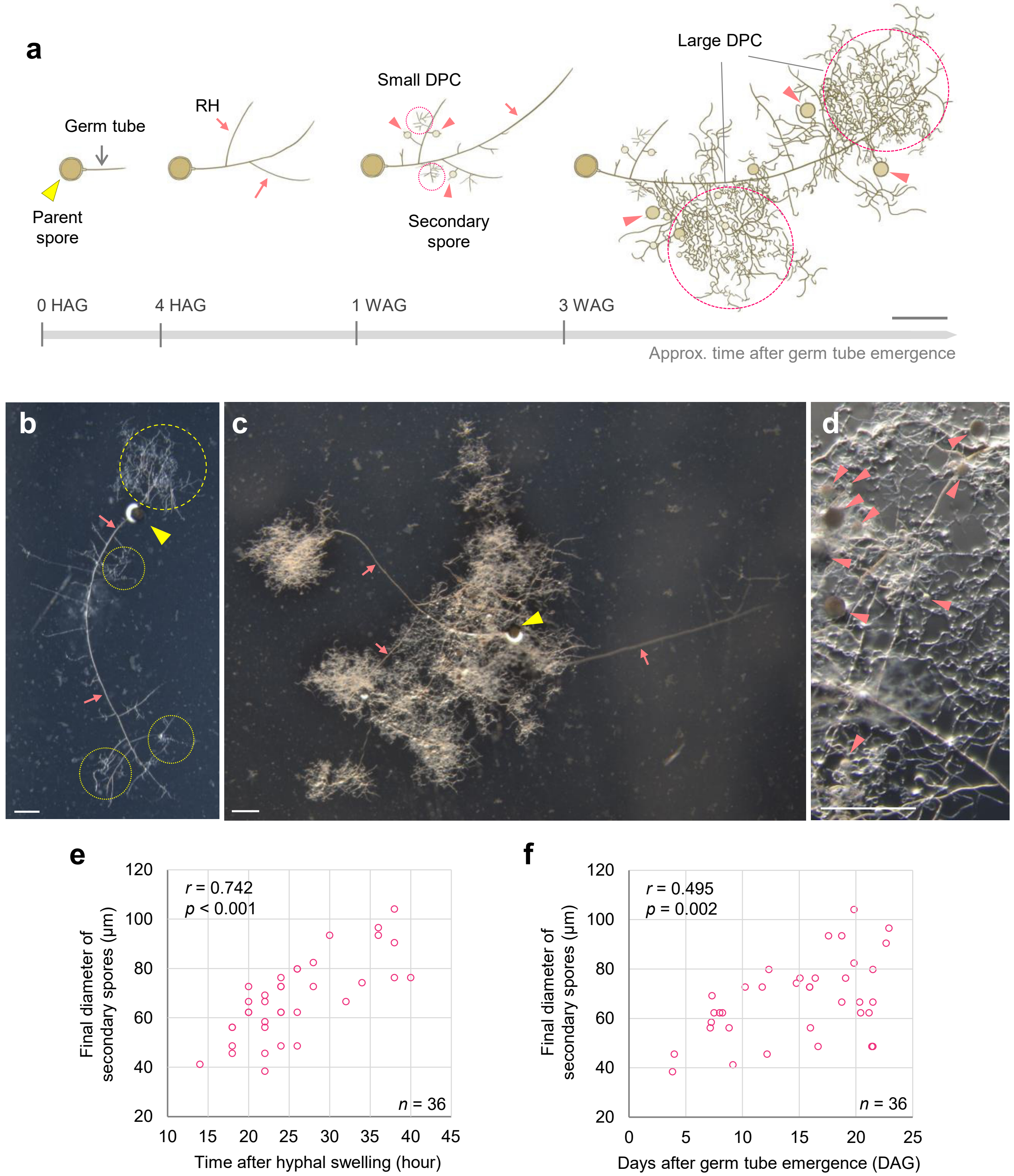


**Supplementary figure 6. Time-lapse analysis of *R. clarus* growth under asymbiotic conditions. a–d**, A schematic diagram (**a**) and representative images (**b–d**) showing *R. clarus* growth in TGM medium. Yellow and pink arrowheads indicate parent and secondary spores. Grey and pink arrows indicate germ tubes and runner hyphae (RH). Dotted and dashed circles indicate small and large DPC (densely packed coils), respectively. HAG, DAG and WAG are hours, days and weeks after germ tube emergence. Bars = 500 µm. **b**, Small DPCs and a developing large DPC at 11 DAG. **c**, Developed large DPCs. **d**, Magnified image of a large DPC. **e–f.** Development of secondary spores in TGM medium. Only focused images were analyzed. **e**, Correlation between the final diameter of secondary spores and time required for the spore development after hyphal swelling. **f**, Correlation between the final diameter of secondary spores and days after germ tube emergence. *r* values are Pearson's correlation coefficient.

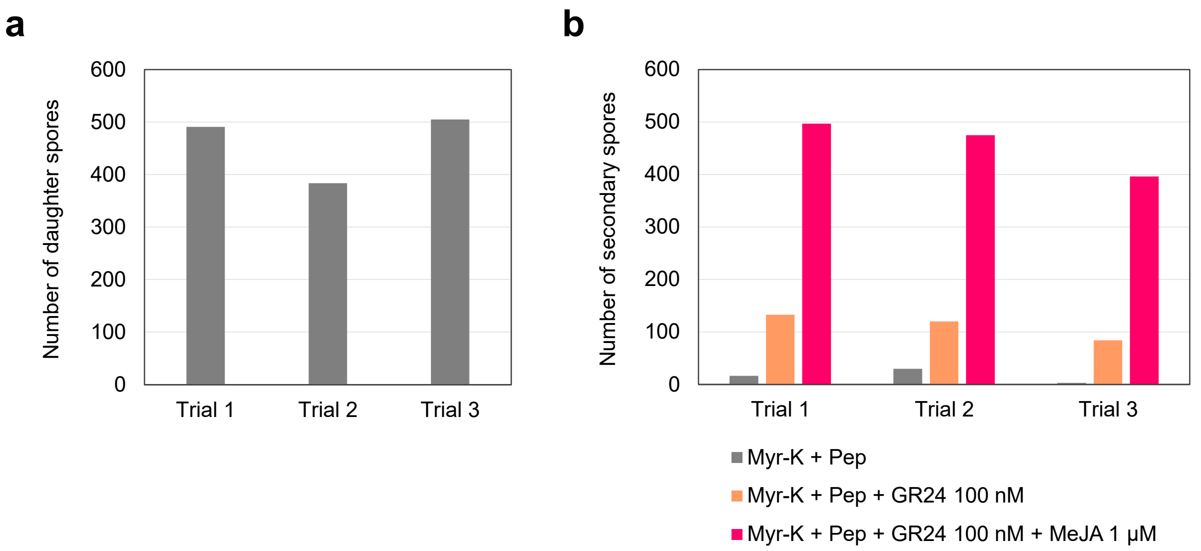


**Supplementary Figure 7. Comparison of the number of spores generated by asymbiotic and *in vitro* monoxenic culture. a**, Numbers of daughter spores generated from a single spore by *in vitro* monoxenic culture at 8 WAI. **b**, Numbers of secondary spores (< 30 µm in diameter) per spore at 8 WAI, which were calculated from the total number of secondary spores produced from 8 parent spores that automatically counted using Ilastik and ImageJ software (see material method).

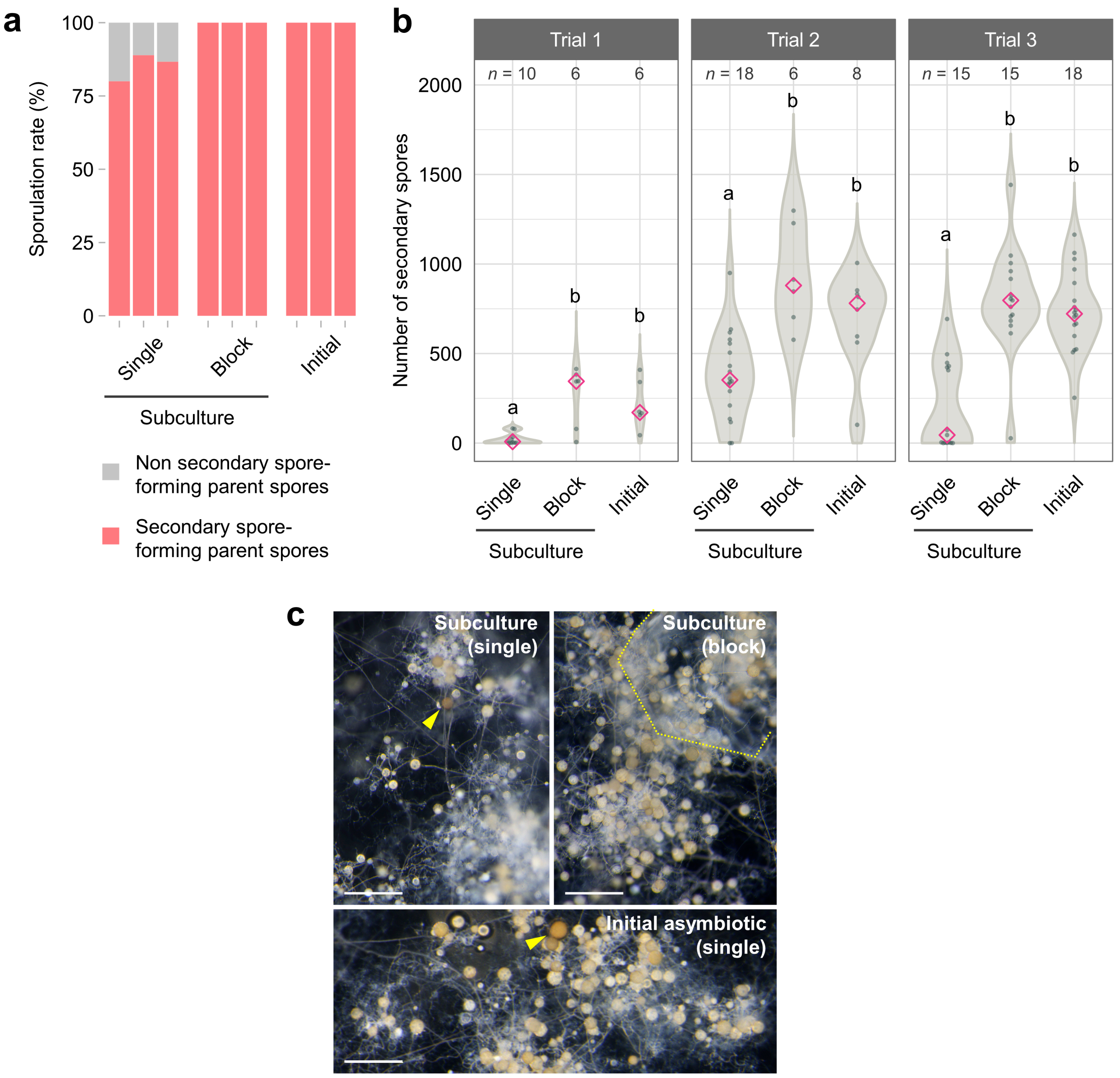


**Supplementary Figure 8. Asymbiotic subculture of *R. clarus*.** The comparison among secondary spore numbers in the initial asymbiotic culture and its subcultures. Single, a single asymbiotically-generated spore was placed on the medium. Block, about 5 mm square gel cut from initial asymbiotic culture medium containing 20–40 spores was placed on the medium. Both initial and subcultures were performed on TGM medium. **a**, Sporulation rate, percentage of secondary spore-forming parent spores relative to germinated patent spores at 6 WAI. **b**, Numbers of secondary spores at 6 WAI. Some block inocula produced too many secondary spores for manual counting, only countable cases (with relatively low number secondary spores) were included. Diamonds indicate medians. Statistical significance was calculated using the Wilcoxon rank-sum test with Bonferroni correction. *Different letters* indicate significant differences ($p < 0.05$). $p$-values are described in Supplementary Table 3. **c**, Secondary spores in the initial asymbiotic culture and its subcultures from single parent spores (arrowheads) or gel block inoculum (dotted polygon) at 8 WAI. Bars = 500 µm.

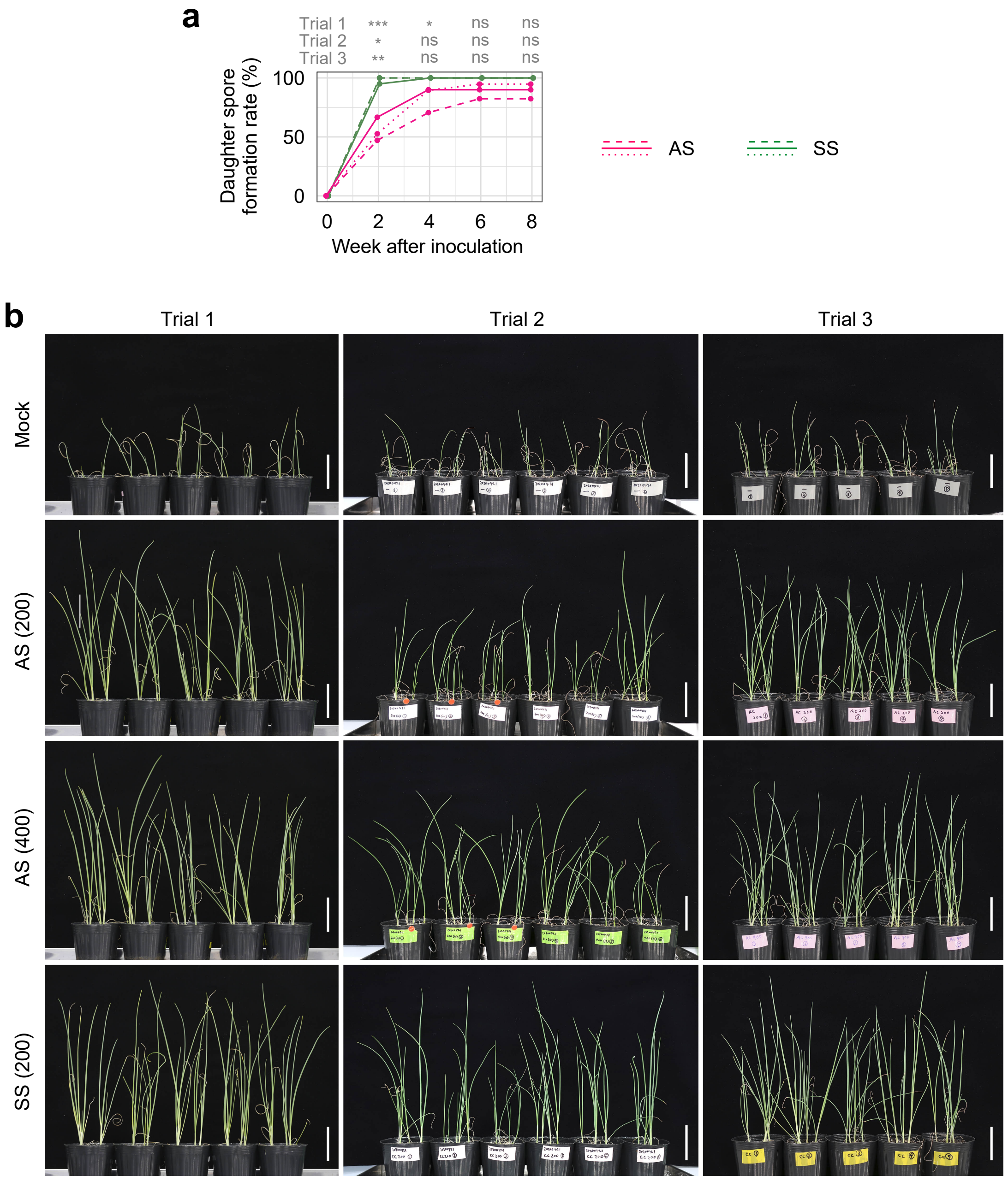


**Supplementary Figure 9. Inoculation tests of *R. clarus* into Welsh onions in pots. a**, Time-course of daughter spore formation rate, percentage of daughter spore-forming spores relative to germinated spores. AS, asymbiotically-generated spores. SS, symbiotically-generated spores. Asterisks above graphs indicate significant difference between treatments in each time point using Fisher's exact test with Bonferroni correction. ***, $p < 0.001$; **, $0.001 \leq p < 0.01$; *, $0.01 \leq p < 0.05$; ns, not significant. *p*-values are described in Supplementary Table 3. **b**, Growth of Welsh onions at 8 weeks after inoculation. Numbers in parentheses are the number of spores inoculated to plants. Bars= 5 cm.

**Supplementary Table 1.** Composition of the modified medium for the asymbiotic culture experiments of *R. clarus* HR1

| | Modified M medium[12] | The modified medium in this study |
|---|---|---|
| **Chemicals** | **Final concentration** | **Final concentration** |
| $MgSO_4 \cdot 7H_2O$ | 731 mg $L^{-1}$ | 731 mg $L^{-1}$ |
| $KNO_3$ | 80 mg $L^{-1}$ | 80 mg $L^{-1}$ |
| KCl | 65 mg $L^{-1}$ | 65 mg $L^{-1}$ |
| $KH_2PO_4$ | 4.8 mg $L^{-1}$ | 4.8 mg $L^{-1}$ |
| $Ca(NO_3) \cdot 4H_2O$ | 288 mg $L^{-1}$ | 288 mg $L^{-1}$ |
| Fe(III)-EDTA | 8 mg $L^{-1}$ | 8 mg $L^{-1}$ |
| $MnCl_2 \cdot 4H_2O$ | 3 mg $L^{-1}$ | 3 mg $L^{-1}$ |
| $ZnSO_4 \cdot 7H_2O$ | 1.3 mg $L^{-1}$ | 1.3 mg $L^{-1}$ |
| $H_3BO_3$ | 1.5 mg $L^{-1}$ | 1.5 mg $L^{-1}$ |
| $CuSO_4 \cdot 5H_2O$ | 0.065 mg $L^{-1}$ | 0.065 mg $L^{-1}$ |
| $Na_2MoO_4 \cdot 2H_2O$ | 0.0012 mg $L^{-1}$ | 0.0012 mg $L^{-1}$ |
| KI | 0.75 mg $L^{-1}$ | 0.75 mg $L^{-1}$ |
| MES (pH 6.5) | 10 mM | 10 mM |
| Sucrose | 10 g $L^{-1}$ | 1 g $L^{-1}$ |
| Glucose | - | 1 g $L^{-1}$ |
| Glycine | 3 mg $L^{-1}$ | 3 mg $L^{-1}$ |
| Pyridoxine-HCl | 0.1 mg $L^{-1}$ | 0.1 mg $L^{-1}$ |
| Nicotinic acid | 0.5 mg $L^{-1}$ | 0.5 mg $L^{-1}$ |
| *myo*-inositol | 50 mg $L^{-1}$ | 50 mg $L^{-1}$ |
| Thiamine-HCl | 0.1 mg $L^{-1}$ | 10 mg $L^{-1}$ |
| Gellan gum (GelGro) | 3.5 g $L^{-1}$ | - |
| Phytagel (Sigma-Aldrich) | - | 3 g $L^{-1}$ |

### Supplementary Table 2. Experimental conditions of asymbiotic cultures and inoculation tests.

**Fig. 1b and Supplementary Fig. 1b**

| Trial | Medium composition (Medium contains chemicals described in Supplementary Table 1 and listed below) | Total number of parent spores | Number of germinated parent spores | Number of secondary spore-forming parent spores | Germination rate (%) | Duration of culture (week) | Number of parent spores per Petri dish | Start of culture | Information of parent spores | | |
|---|---|---|---|---|---|---|---|---|---|---|---|
| | | | | | | | | | Culture method | Start of culture | Duration of culture (month) |
| 1 | Mock | 20 | 19 | 0 | 95.0 | 6 | 10 | January, 2020 | Monoxenic culture | August, 2019 | 5 |
| | Potassium myristate (Myr-K) 100 µM | 20 | 20 | 3 | 100.0 | 6 | 10 | | | | |
| | Potassium palmitate 100 µM | 20 | 16 | 0 | 80.0 | 6 | 10 | | | | |
| | Potassium palmitoleate 100 µM | 20 | 19 | 0 | 95.0 | 6 | 10 | | | | |
| | 2OH-TDA 100 µM | 20 | 18 | 0 | 90.0 | 6 | 10 | | | | |
| 2 | Mock | 20 | 18 | 0 | 90.0 | 6 | 10 | January, 2020 | Monoxenic culture | August, 2019 | 5 |
| | Potassium myristate (Myr-K) 100 µM | 20 | 20 | 0 | 100.0 | 6 | 10 | | | | |
| | Potassium palmitate 100 µM | 20 | 16 | 0 | 80.0 | 6 | 10 | | | | |
| | Potassium palmitoleate 100 µM | 20 | 16 | 0 | 80.0 | 6 | 10 | | | | |
| | 2OH-TDA 100 µM | 20 | 17 | 0 | 85.0 | 6 | 10 | | | | |
| 3 | Mock | 20 | 19 | 0 | 95.0 | 6 | 10 | January, 2020 | Monoxenic culture | August, 2019 | 5 |
| | Potassium myristate (Myr-K) 100 µM | 20 | 20 | 0 | 100.0 | 6 | 10 | | | | |
| | Potassium palmitate 100 µM | 20 | 20 | 0 | 100.0 | 6 | 10 | | | | |
| | Potassium palmitoleate 100 µM | 20 | 18 | 1 | 90.0 | 6 | 10 | | | | |
| | 2OH-TDA 100 µM | 20 | 20 | 0 | 100.0 | 6 | 10 | | | | |

**Fig. 1c, d (trial 2) and Supplementary Fig. 2a (trial 1 and 3)**

| Trial | Medium composition (Medium contains chemicals described in Supplementary Table 1 and listed below) | Total number of parent spores | Number of germinated parent spores | Number of secondary spore-forming parent spores | Germination rate (%) | Duration of culture (week) | Number of parent spores per Petri dish | Start of culture | Information of parent spores | | |
|---|---|---|---|---|---|---|---|---|---|---|---|
| | | | | | | | | | Culture method | Start of culture | Duration of culture (month) |
| 1 | Myr-K 100 µM | 20 | 20 | 0 | 100.0 | 6 | 5 | November, 2019 | Monoxenic culture | April, 2019 | 7 |
| | Myr-K 100 µM + Malt extract 0.2 mg $L^{-1}$ | 20 | 19 | 5 | 95.0 | 6 | 5 | | | | |
| | Myr-K 100 µM + SC dropout 0.2 mg $L^{-1}$ | 20 | 20 | 9 | 100.0 | 6 | 5 | | | | |
| | Myr-K 100 µM + Yeast extract 0.2 mg $L^{-1}$ | 20 | 19 | 18 | 95.0 | 6 | 5 | | | | |
| | Myr-K 100 µM + Peptone 0.2 mg $L^{-1}$ | 20 | 18 | 18 | 90.0 | 6 | 5 | | | | |
| 2 | Myr-K 100 µM | 20 | 19 | 5 | 95.0 | 6 | 5 | November, 2019 | Monoxenic culture | April, 2019 | 7 |
| | Myr-K 100 µM + Malt extract 0.2 mg $L^{-1}$ | 20 | 19 | 1 | 95.0 | 6 | 5 | | | | |
| | Myr-K 100 µM + SC dropout 0.2 mg $L^{-1}$ | 20 | 18 | 15 | 90.0 | 6 | 5 | | | | |
| | Myr-K 100 µM + Yeast extract 0.2 mg $L^{-1}$ | 20 | 19 | 17 | 95.0 | 6 | 5 | | | | |
| | Myr-K 100 µM + Peptone 0.2 mg $L^{-1}$ | 20 | 18 | 16 | 90.0 | 6 | 5 | | | | |
| 3 | Myr-K 100 µM | 20 | 19 | 4 | 95.0 | 6 | 5 | November, 2019 | Monoxenic culture | May, 2019 | 6 |
| | Myr-K 100 µM + Malt extract 0.2 mg $L^{-1}$ | 20 | 20 | 9 | 100.0 | 6 | 5 | | | | |
| | Myr-K 100 µM + SC dropout 0.2 mg $L^{-1}$ | 20 | 18 | 5 | 90.0 | 6 | 5 | | | | |
| | Myr-K 100 µM + Yeast extract 0.2 mg $L^{-1}$ | 20 | 18 | 14 | 90.0 | 6 | 5 | | | | |
| | Myr-K 100 µM + Peptone 0.2 mg $L^{-1}$ | 20 | 19 | 9 | 95.0 | 6 | 5 | | | | |

| Trial | Medium composition (Medium contains chemicals described in Supplementary Table 1 and listed below) | Total number of parent spores | Number of germinated parent spores | Number of secondary spore-forming parent spores | Germination rate (%) | Duration of culture (week) | Number of parent spores per Petri dish | Start of culture | Information of parent spores | | |
|---|---|---|---|---|---|---|---|---|---|---|---|
| | | | | | | | | | Culture method | Start of culture | Duration of culture (month) |
| **Supplementary Fig. 2c** | | | | | | | | | | | |
| 1 | Myr-K 100 µM + Peptone 0.2 mg $L^{-1}$ | 20 | 20 | 17 | 100.0 | 6 | 5 | December, 2019 | Monoxenic culture | April, 2019 | 7 |
| | Myr-K 100 µM + Peptone 1 mg $L^{-1}$ | 20 | 20 | 16 | 100.0 | 6 | 5 | | | | |
| | Myr-K 500 µM + Peptone 0.2 mg $L^{-1}$ | 20 | 20 | 19 | 100.0 | 6 | 5 | | | | |
| | Myr-K 500 µM + Peptone 1 mg $L^{-1}$ | 20 | 20 | 13 | 100.0 | 6 | 5 | | | | |
| 2 | Myr-K 100 µM + Peptone 0.2 mg $L^{-1}$ | 20 | 20 | 15 | 100.0 | 6 | 5 | December, 2019 | Monoxenic culture | April, 2019 | 7 |
| | Myr-K 100 µM + Peptone 1 mg $L^{-1}$ | 20 | 19 | 15 | 95.0 | 6 | 5 | | | | |
| | Myr-K 500 µM + Peptone 0.2 mg $L^{-1}$ | 20 | 20 | 18 | 100.0 | 6 | 5 | | | | |
| | Myr-K 500 µM + Peptone 1 mg $L^{-1}$ | 20 | 20 | 9 | 100.0 | 6 | 5 | | | | |
| 3 | Myr-K 100 µM + Peptone 0.2 mg $L^{-1}$ | 20 | 20 | 10 | 100.0 | 6 | 5 | December, 2019 | Monoxenic culture | May, 2019 | 7 |
| | Myr-K 100 µM + Peptone 1 mg $L^{-1}$ | 15 | 15 | 11 | 100.0 | 6 | 5 | | | | |
| | Myr-K 500 µM + Peptone 0.2 mg $L^{-1}$ | 20 | 20 | 9 | 100.0 | 6 | 5 | | | | |
| | Myr-K 500 µM + Peptone 1 mg $L^{-1}$ | 20 | 20 | 9 | 100.0 | 6 | 5 | | | | |
| **Fig. 2b (trial 1) and Supplementary Fig. 2a (trial 2 and 3)** | | | | | | | | | | | |
| 1 | Myr-K 500 µM + Peptone 1 mg $L^{-1}$ | 16 | 12 | 8 | 75.0 | 6 | 4 | October, 2019 | Monoxenic culture | April, 2019 | 6 |
| | Myr-K 500 µM + Peptone 1 mg $L^{-1}$ + GR24 10 nM | 16 | 15 | 15 | 93.8 | 6 | 4 | | | | |
| | Myr-K 500 µM + Peptone 1 mg $L^{-1}$ + GR24 100 nM | 16 | 14 | 14 | 87.5 | 6 | 4 | | | | |
| | Myr-K 500 µM + Peptone 1 mg $L^{-1}$ + GR24 1 µM | 16 | 13 | 13 | 81.3 | 6 | 4 | | | | |
| 2 | Myr-K 500 µM + Peptone 1 mg $L^{-1}$ | 16 | 13 | 10 | 81.3 | 6 | 4 | October, 2019 | Monoxenic culture | April, 2019 | 6 |
| | Myr-K 500 µM + Peptone 1 mg $L^{-1}$ + GR24 10 nM | 12 | 9 | 9 | 75.0 | 6 | 4 | | | | |
| | Myr-K 500 µM + Peptone 1 mg $L^{-1}$ + GR24 100 nM | 16 | 14 | 14 | 87.5 | 6 | 4 | | | | |
| | Myr-K 500 µM + Peptone 1 mg $L^{-1}$ + GR24 1 µM | 16 | 13 | 13 | 81.3 | 6 | 4 | | | | |
| 3 | Myr-K 500 µM + Peptone 1 mg $L^{-1}$ | 20 | 19 | 6 | 95.0 | 6 | 4 | December, 2019 | Monoxenic culture | April, 2019 | 8 |
| | Myr-K 500 µM + Peptone 1 mg $L^{-1}$ + GR24 10 nM | 20 | 19 | 19 | 95.0 | 6 | 4 | | | | |
| | Myr-K 500 µM + Peptone 1 mg $L^{-1}$ + GR24 100 nM | 20 | 18 | 15 | 90.0 | 6 | 4 | | | | |
| | Myr-K 500 µM + Peptone 1 mg $L^{-1}$ + GR24 1 µM | 20 | 18 | 18 | 90.0 | 6 | 4 | | | | |

**Fig. 2c, d**

| Trial | Medium composition (Medium contains chemicals described in Supplementary Table 1 and listed below) | Total number of parent spores | Number of germinated parent spores | Number of secondary spore-forming parent spores | Germination rate (%) | Duration of culture (days) | Number of parent spores per Petri dish | Start of culture | Information of parent spores | | |
|---|---|---|---|---|---|---|---|---|---|---|---|
| | | | | | | | | | Culture method | Start of culture | Duration of culture (month) |
| 1 | Myr-K 500 µM + Peptone 1 mg $L^{-1}$ | 41 | 13 | 0 | 31.7 | 5 | 20–21 | October, 2019 | Monoxenic culture | April, 2019 | 6 |
| | Myr-K 500 µM + Peptone 1 mg $L^{-1}$ | 41 | 15 | 0 | 36.6 | 8 | 20–21 | | | | |
| | Myr-K 500 µM + Peptone 1 mg $L^{-1}$ | 41 | 38 | 2 | 92.7 | 14 | 20–21 | | | | |
| | Myr-K 500 µM + Peptone 1 mg $L^{-1}$ + GR24 100 nM | 48 | 39 | 0 | 81.3 | 5 | 24 | | | | |
| | Myr-K 500 µM + Peptone 1 mg $L^{-1}$ + GR24 100 nM | 48 | 41 | 11 | 85.4 | 8 | 24 | | | | |
| | Myr-K 500 µM + Peptone 1 mg $L^{-1}$ + GR24 100 nM | 48 | 44 | 39 | 91.7 | 14 | 24 | | | | |
| 2 | Myr-K 500 µM + Peptone 1 mg $L^{-1}$ | 40 | 2 | 0 | 5.0 | 5 | 20 | December, 2019 | Monoxenic culture | August, 2019 | 5 |
| | Myr-K 500 µM + Peptone 1 mg $L^{-1}$ | 40 | 8 | 0 | 20.0 | 8 | 20 | | | | |
| | Myr-K 500 µM + Peptone 1 mg $L^{-1}$ | 40 | 31 | 2 | 77.5 | 14 | 20 | | | | |
| | Myr-K 500 µM + Peptone 1 mg $L^{-1}$ + GR24 100 nM | 40 | 38 | 0 | 95.0 | 5 | 20 | | | | |
| | Myr-K 500 µM + Peptone 1 mg $L^{-1}$ + GR24 100 nM | 40 | 39 | 11 | 97.5 | 8 | 20 | | | | |
| | Myr-K 500 µM + Peptone 1 mg $L^{-1}$ + GR24 100 nM | 40 | 39 | 36 | 97.5 | 14 | 20 | | | | |
| 3 | Myr-K 500 µM + Peptone 1 mg $L^{-1}$ | 40 | 1 | 0 | 2.5 | 5 | 20 | December, 2019 | Monoxenic culture | August, 2019 | 5 |
| | Myr-K 500 µM + Peptone 1 mg $L^{-1}$ | 40 | 5 | 0 | 12.5 | 8 | 20 | | | | |
| | Myr-K 500 µM + Peptone 1 mg $L^{-1}$ | 40 | 34 | 0 | 85.0 | 14 | 20 | | | | |
| | Myr-K 500 µM + Peptone 1 mg $L^{-1}$ + GR24 100 nM | 40 | 28 | 0 | 70.0 | 5 | 20 | | | | |
| | Myr-K 500 µM + Peptone 1 mg $L^{-1}$ + GR24 100 nM | 40 | 35 | 4 | 87.5 | 8 | 20 | | | | |
| | Myr-K 500 µM + Peptone 1 mg $L^{-1}$ + GR24 100 nM | 40 | 36 | 30 | 90.0 | 14 | 20 | | | | |

**Fig. 2e (trial 1) and Supplementary Fig. 4 (trial 2 and 3)**

| Trial | Medium composition (Medium contains chemicals described in Supplementary Table 1 and listed below) | Total number of parent spores | Number of germinated parent spores | Number of secondary spore-forming parent spores | Germination rate (%) | Duration of culture (week) | Number of parent spores per Petri dish | Start of culture | Information of parent spores | | |
|---|---|---|---|---|---|---|---|---|---|---|---|
| | | | | | | | | | Culture method | Start of culture | Duration of culture (month) |
| 1 | T medium (Myr-K 500 µM + Peptone 1 mg $L^{-1}$) | 16 | 13 | 11 | 81.3 | 6 | 4 | October, 2019 | Monoxenic culture | April, 2019 | 6 |
| | Myr-K 500 µM + Peptone 1 mg $L^{-1}$ + MeJA 1 µM | 16 | 15 | 15 | 93.8 | 6 | 4 | | | | |
| | Myr-K 500 µM + Peptone 1 mg $L^{-1}$ + GR24 100 nM | 17 | 15 | 15 | 88.2 | 6 | 4 | | | | |
| | Myr-K 500 µM + Peptone 1 mg $L^{-1}$ + GR24 100 nM + MeJA 0.1 µM | 16 | 14 | 14 | 87.5 | 6 | 4 | | | | |
| | Myr-K 500 µM + Peptone 1 mg $L^{-1}$ + GR24 100 nM + MeJA 1 µM | 36 | 31 | 31 | 86.1 | 6 | 4 | | | | |
| | Myr-K 500 µM + Peptone 1 mg $L^{-1}$ + GR24 100 nM + MeJA 10 µM | 16 | 12 | 12 | 75.0 | 6 | 4 | | | | |
| 2 | T medium (Myr-K 500 µM + Peptone 1 mg $L^{-1}$) | 16 | 14 | 14 | 87.5 | 6 | 4 | October, 2019 | Monoxenic culture | April, 2019 | 6 |
| | Myr-K 500 µM + Peptone 1 mg $L^{-1}$ + MeJA 1 µM | 12 | 11 | 11 | 91.7 | 6 | 4 | | | | |
| | Myr-K 500 µM + Peptone 1 mg $L^{-1}$ + GR24 100 nM | 16 | 14 | 14 | 87.5 | 6 | 4 | | | | |
| | Myr-K 500 µM + Peptone 1 mg $L^{-1}$ + GR24 100 nM + MeJA 0.1 µM | 16 | 13 | 13 | 81.3 | 6 | 4 | | | | |
| | Myr-K 500 µM + Peptone 1 mg $L^{-1}$ + GR24 100 nM + MeJA 1 µM | 32 | 29 | 29 | 90.6 | 6 | 4 | | | | |
| | Myr-K 500 µM + Peptone 1 mg $L^{-1}$ + GR24 100 nM + MeJA 10 µM | 16 | 14 | 14 | 87.5 | 6 | 4 | | | | |
| 3 | T medium (Myr-K 500 µM + Peptone 1 mg $L^{-1}$) | 20 | 19 | 5 | 95.0 | 6 | 4 | December, 2019 | Monoxenic culture | August, 2019 | 4 |
| | Myr-K 500 µM + Peptone 1 mg $L^{-1}$ + MeJA 1 µM | 20 | 20 | 9 | 100.0 | 6 | 4 | | | | |
| | Myr-K 500 µM + Peptone 1 mg $L^{-1}$ + GR24 100 nM | 20 | 19 | 18 | 95.0 | 6 | 4 | | | | |
| | Myr-K 500 µM + Peptone 1 mg $L^{-1}$ + GR24 100 nM + MeJA 0.1 µM | 20 | 20 | 20 | 100.0 | 6 | 4 | | | | |
| | Myr-K 500 µM + Peptone 1 mg $L^{-1}$ + GR24 100 nM + MeJA 1 µM | 20 | 20 | 20 | 100.0 | 6 | 4 | | | | |
| | Myr-K 500 µM + Peptone 1 mg $L^{-1}$ + GR24 100 nM + MeJA 10 µM | 20 | 19 | 19 | 95.0 | 6 | 4 | | | | |

| Trial | Experimental conditions (Medium contains chemicals described in Supplementary Table 1 and listed below. All experiments were performed on TGM medium) | | Total number of parent spores | Number of germinated parent spores | Number of secondary spore-forming parent spores | Germination rate (%) | Duration of culture (week) | Number of parent spores per Petri dish | Start of culture | Information of parent spores | | |
|---|---|---|---|---|---|---|---|---|---|---|---|---|
| | | | | | | | | | | Culture method | Start of culture | Duration of culture (month) |
| **Supplementary Fig. 5** | | | | | | | | | | | | |
| 1 | *R. irregularis* asymbiotic culture | | 15 | 12 | 8 | 80.0 | 8 | 5 | February, 2020 | Purchased from Premier Tech at May, 2017. Lot number: 10889843 | | |
| 2 | *R. irregularis* asymbiotic culture | | 20 | 11 | 6 | 55.0 | 8 | 5 | July, 2020 | Purchased from Premier Tech at November, 2018. Lot number: 11426795 | | |
| 3 | *R. irregularis* asymbiotic culture | | 20 | 17 | 12 | 85.0 | 8 | 5 | July, 2020 | Purchased from Premier Tech at November, 2018. Lot number: 11426795 | | |
| **Supplementary Fig. 8** | | | | | | | | | | | | |
| 1 | Subculture (2nd asymbiotic culture) | single spore placed | 10 | 10 | 8 | 100.0 | 8 | 5 | January, 2020 | Asymbiotic culture | September, 2019 | 4 |
| | | block placed | 6 | 6 | 6 | 100.0 | 8 | 3 | | | | |
| | Initial asymbiotic culture (single spore placed) | | 6 | 6 | 6 | 100.0 | 8 | 3 | | Monoxenic culture | August, 2019 | 5 |
| 2 | Subculture (2nd asymbiotic culture) | single spore placed | 20 | 18 | 16 | 90.0 | 8 | 5 | April, 2020 | Asymbiotic culture | Nobember, 2019 | 5 |
| | | block placed | 12 | 12 | 12 | 100.0 | 8 | 3 | | | | |
| | Initial asymbiotic culture (single spore placed) | | 8 | 8 | 8 | 100.0 | 8 | 4 | | Monoxenic culture | November, 2019 | 5 |
| 3 | Subculture (2nd asymbiotic culture) | single spore placed | 20 | 15 | 13 | 75.0 | 8 | 4 | July, 2020 | Asymbiotic culture | February, 2020 | 5 |
| | | block placed | 15 | 15 | 15 | 100.0 | 8 | 3 | | | | |
| | Initial asymbiotic culture (single spore placed) | | 20 | 18 | 18 | 90.0 | 8 | 4 | | Monoxenic culture | January, 2020 | 6 |

**Fig. 3a, b**

| Trial | Spores inoculated to plants | Number of spores inoculated to plants | Number of germinated spores | | | | Number of spores formed daughter spores | | | | Information of spores inoculated to plants | |
|---|---|---|---|---|---|---|---|---|---|---|---|---|
| | | | 2 WAI | 4 WAI | 6 WAI | 8 WAI | 2 WAI | 4 WAI | 6 WAI | 8 WAI | Start of culture | Culture period |
| 1 | Asymbiotically generated spores (AS) | 20 | 17 | 17 | 17 | 17 | 8 | 12 | 14 | 14 | July, 2019 | 4 months |
| | Symbiotically generated spores (SS) | 21 | 20 | 20 | 20 | 20 | 20 | 20 | 20 | 20 | April, 2019 | 8 months |
| 2 | AS | 20 | 18 | 20 | 20 | 20 | 12 | 18 | 18 | 18 | July, 2019 | 4 months |
| | SS | 21 | 20 | 21 | 21 | 21 | 19 | 21 | 21 | 21 | April, 2019 | 8 months |
| 3 | AS | 20 | 19 | 19 | 19 | 19 | 10 | 17 | 18 | 18 | July, 2019 | 4 months |
| | SS | 20 | 18 | 20 | 20 | 20 | 18 | 20 | 20 | 20 | May, 2019 | 7 months |

| Trial | Spores inoculated to plants | Germination rate (%) | | | | Sporulation rate (%) | | | | | | | |
|---|---|---|---|---|---|---|---|---|---|---|---|---|---|
| | | | | | | Per spores inoculated to plants | | | | Per germinated spores | | | |
| | | 2 WAI | 4 WAI | 6 WAI | 8 WAI | 2 WAI | 4 WAI | 6 WAI | 8 WAI | 2 WAI | 4 WAI | 6 WAI | 8 WAI |
| 1 | AS | 85.0 | 85.0 | 85.0 | 85.0 | 40.0 | 60.0 | 70.0 | 70.0 | 47.1 | 70.6 | 82.4 | 82.4 |
| | SS | 95.2 | 95.2 | 95.2 | 95.2 | 95.2 | 95.2 | 95.2 | 95.2 | 100.0 | 100.0 | 100.0 | 100.0 |
| 2 | AS | 90.0 | 100.0 | 100.0 | 100.0 | 60.0 | 90.0 | 90.0 | 90.0 | 66.7 | 90.0 | 90.0 | 90.0 |
| | SS | 95.2 | 100.0 | 100.0 | 100.0 | 90.5 | 100.0 | 100.0 | 100.0 | 95.0 | 100.0 | 100.0 | 100.0 |
| 3 | AS | 95.0 | 95.0 | 95.0 | 95.0 | 50.0 | 85.0 | 90.0 | 90.0 | 52.6 | 89.5 | 94.7 | 94.7 |
| | SS | 90.0 | 100.0 | 100.0 | 100.0 | 90.0 | 100.0 | 100.0 | 100.0 | 100.0 | 100.0 | 100.0 | 100.0 |

**Fig. 3g**

| Trial | Spores inoculated to plants | Number of spores inoculated to pots | Number of plants per pot | Number of pots | Start of culture | Information of spores inoculated to plants | |
|---|---|---|---|---|---|---|---|
| | | | | | | Start of culture | Duration of culture |
| 1 | Mock | - | 4 | 5 | January, 2020 | | |
| | Asymbiotically generated spores (AS) | 200 | 4 | 5 | | July or September, 2019 | 4–6 months |
| | | 400 | 4 | 5 | | | |
| | Symbiotically generated spores (SS) | 200 | 4 | 5 | | July or September, 2019 | 4–6 months |
| 2 | Mock | - | 4 | 5 | April, 2020 | | |
| | AS | 200 | 4 | 5 | | October or November, 2019 | 5–6 months |
| | | 400 | 4 | 4 | | | |
| | SS | 200 | 4 | 4 | | November, 2019 | 5 months |
| 3 | Mock | - | 4 | 4 | June, 2020 | | |
| | AS | 200 | 4 | 4 | | December, 2019 | 6 months |
| | | 400 | 4 | 4 | | | |
| | SS | 200 | 4 | 4 | | December, 2019 | 6 months |

**Supplementary Table 3.** *p*-values and effect size of all statistical analysis. ***, $p < 0.001$. **, $0.001 \le p < 0.01$. *, $0.01 \le p < 0.05$. ns, not significant.

**Supplementary Fig. 1b**

| Condition | Maximum feret diameter | | | | | |
|---|---|---|---|---|---|---|
| | Trial 1 | | Trial 2 | | Trial 3 | |
| | *p*-value | Significance | *p*-value | Significance | *p*-value | Significance |
| Mock – C14:0-K (Myr-K) | < 0.001 | *** | 0.002 | ** | < 0.001 | *** |
| Mock – C16:0-K | 1.000 | ns | 1.000 | ns | 1.000 | ns |
| Mock – C16:1-K | < 0.001 | *** | < 0.001 | *** | < 0.001 | *** |
| Mock – 2OH-TDA | 1.000 | ns | 1.000 | ns | 1.000 | ns |
| C14:0-K (Myr-K) – C16:0-K | 0.007 | ** | 0.006 | ** | < 0.001 | *** |
| C14:0-K (Myr-K) – C16:1-K | < 0.001 | *** | < 0.001 | *** | < 0.001 | *** |
| C14:0-K (Myr-K) – 2OH-TDA | < 0.001 | *** | 0.006 | ** | < 0.001 | *** |
| C16:0-K – C16:1-K | < 0.001 | *** | < 0.001 | *** | < 0.001 | *** |
| C16:0-K – 2OH-TDA | 1.000 | ns | 1.000 | ns | 1.000 | ns |
| C16:1-K – 2OH-TDA | < 0.001 | *** | 0.001 | ** | < 0.001 | *** |

| Condition | Minimum feret diameter | | | | | |
|---|---|---|---|---|---|---|
| | Trial 1 | | Trial 2 | | Trial 3 | |
| | *p*-value | Significance | *p*-value | Significance | *p*-value | Significance |
| Mock – C14:0-K (Myr-K) | 0.002 | ** | 0.001 | ** | < 0.001 | *** |
| Mock – C16:0-K | 0.881 | ns | 1.000 | ns | 1.000 | ns |
| Mock – C16:1-K | 0.002 | ** | < 0.001 | *** | 0.977 | ns |
| Mock – 2OH-TDA | 0.010 | ** | 0.004 | ** | 0.891 | ns |
| C14:0-K (Myr-K) – C16:0-K | < 0.001 | *** | 0.006 | ** | < 0.001 | *** |
| C14:0-K (Myr-K) – C16:1-K | < 0.001 | *** | < 0.001 | *** | < 0.001 | *** |
| C14:0-K (Myr-K) – 2OH-TDA | < 0.001 | *** | < 0.001 | *** | < 0.001 | *** |
| C16:0-K – C16:1-K | 0.765 | ns | < 0.001 | *** | 0.294 | ns |
| C16:0-K – 2OH-TDA | 1.000 | ns | 0.002 | ** | 0.256 | ns |
| C16:1-K – 2OH-TDA | 1.000 | ns | 1.000 | ns | 1.000 | ns |

**Fig. 1c**

| Trial | Condition | *p*-value | | | |
|---|---|---|---|---|---|
| | | Mock | Malt | SC | Yeast |
| 1 | Malt | 0.202 (ns) | - | - | - |
| | SC | 0.012 (*) | 1.000 (ns) | - | - |
| | Yeast | < 0.001 (***) | < 0.001 (***) | 0.013 (*) | - |
| | Pep | < 0.001 (***) | < 0.001 (***) | 0.002 (**) | 1.000 (ns) |
| 2 | Malt | 1.000 (ns) | - | - | - |
| | SC | 0.008 (**) | < 0.001 (***) | - | - |
| | Yeast | 0.002 (**) | < 0.001 (***) | 1.000 (ns) | - |
| | Pep | 0.002 (**) | < 0.001 (***) | 1.000 (ns) | 1.000 (ns) |
| 3 | Malt | 1.000 (ns) | - | - | - |
| | SC | 1.000 (ns) | 1.000 (ns) | - | - |
| | Yeast | 0.009 (**) | 0.521 (ns) | 0.067 (ns) | - |
| | Pep | 1.000 (ns) | 1.000 (ns) | 1.000 (ns) | 0.911 (ns) |

**Fig. 1d**

| Condition | Trial 2 | | |
|---|---|---|---|
| | *p*-value | Hedge's *g* | Significance |
| Mock – Malt | 1.000 | 0.390 | ns |
| SC – Mock | < 0.001 | 1.352 | *** |
| Yeast – Mock | < 0.001 | 1.036 | *** |
| Pep – Mock | < 0.001 | 1.254 | *** |
| SC – Malt | < 0.001 | 1.430 | *** |
| Pep – Malt | < 0.001 | 1.270 | *** |
| Yeast – SC | 0.940 | 0.500 | ns |
| SC – Pep | 0.514 | 1.000 | ns |
| Yeast – Malt | < 0.001 | 1.064 | *** |
| Yeast – Pep | 1.000 | 0.710 | ns |

**Supplementary Fig. 2a**

| Condition | Trial 1 | | | Trial 3 | | |
|---|---|---|---|---|---|---|
| | *p*-value | Hedge's *g* | Significance | *p*-value | Hedge's *g* | Significance |
| Mock – Malt | 0.095 | 0.760 | ns | 0.691 | 0.752 | ns |
| SC – Mock | 0.009 | 0.591 | ** | 1.000 | 0.305 | ns |
| Yeast – Mock | < 0.001 | 1.343 | *** | 0.001 | 1.062 | ** |
| Pep – Mock | < 0.001 | 1.452 | *** | 1.000 | 0.804 | ns |
| SC – Malt | 1.000 | 0.446 | ns | 1.000 | 0.539 | ns |
| Pep – Malt | < 0.001 | 1.421 | *** | 1.000 | 0.629 | ns |
| Yeast – SC | < 0.001 | 1.187 | *** | 0.015 | 0.949 | * |
| SC – Pep | < 0.001 | 1.403 | *** | 1.000 | 0.752 | ns |
| Yeast – Malt | < 0.001 | 1.295 | *** | 0.369 | 0.642 | ns |
| Yeast – Pep | 0.288 | 0.918 | ns | 0.586 | 0.280 | ns |

**Fig. 2a**

| Trial | Conditions | *p*-value | | |
|---|---|---|---|---|
| | | Mock | GR24 10nM | GR24 100nM |
| 1 | GR24 10 nM | 0.169 (ns) | - | - |
| | GR24 100 nM | 0.199 (ns) | 1.000 (ns) | - |
| | GR24 1 μM | 0.235 (ns) | 1.000 (ns) | 1.000 (ns) |
| 2 | GR24 10 nM | 1.000 (ns) | - | - |
| | GR24 100 nM | 0.587 (ns) | 1.000 (ns) | - |
| | GR24 1 μM | 1.000 (ns) | 1.000 (ns) | 1.000 (ns) |
| 3 | GR24 10 nM | < 0.001 (***) | - | - |
| | GR24 100 nM | 0.015 (*) | 0.630 (ns) | - |
| | GR24 1 μM | < 0.001(***) | 1.000 (ns) | 1.000 (ns) |

**Fig. 2b**

| Condition | Trial 1 | | |
|---|---|---|---|
| | *p*-value | Hedge's *g* | Significance |
| Mock – GR24 10 nM | < 0.001 | 2.661 | *** |
| Mock – GR24 100 nM | < 0.001 | 1.888 | *** |
| Mock – GR24 1 µM | < 0.001 | 2.028 | *** |
| GR24 100 nM – GR24 10 nM | 1.000 | 0.404 | ns |
| GR24 10 nM – GR24 1 µM | 0.144 | 0.996 | ns |
| GR24 100 nM – GR24 1 µM | 1.000 | 0.592 | ns |

**Supplementary Fig. 3a**

| Condition | Trial 2 | | | Trial 3 | | |
|---|---|---|---|---|---|---|
| | *p*-value | Hedge's *g* | Significance | *p*-value | Hedge's *g* | Significance |
| Mock – GR24 10 nM | 0.017 | 1.837 | * | < 0.001 | 2.653 | *** |
| Mock – GR24 100 nM | < 0.001 | 2.249 | *** | < 0.001 | 1.527 | *** |
| Mock – GR24 1 µM | < 0.001 | 1.718 | *** | < 0.001 | 3.115 | *** |
| GR24 100 nM – GR24 10 nM | 0.379 | 0.879 | ns | 1.000 | 0.178 | ns |
| GR24 10 nM – GR24 1 µM | 0.165 | 0.948 | ns | < 0.001 | 1.454 | *** |
| GR24 100 nM – GR24 1 µM | 1.000 | 0.469 | ns | 0.001 | 1.391 | ** |

**Fig. 2c**

| Condition | Duration of culture (days) | Trial 1 | | Trial 2 | | Trial 3 | |
|---|---|---|---|---|---|---|---|
| | | *p*-value | Significance | *p*-value | Significance | *p*-value | Significance |
| GR24 - – GR24 + | 5 | < 0.001 | *** | < 0.001 | *** | < 0.001 | *** |
| | 8 | < 0.001 | *** | < 0.001 | *** | < 0.001 | *** |
| | 14 | 1.000 | ns | 0.014 | * | 0.737 | ns |

**Fig. 2d**

| Condition | Duration of culture (days) | Trial 1 | | Trial 2 | | Trial 3 | |
|---|---|---|---|---|---|---|---|
| | | *p*-value | Significance | *p*-value | Significance | *p*-value | Significance |
| GR24 - – GR24 + | 5 | 1.000 | ns | 1.000 | ns | 1.000 | ns |
| | 8 | 0.026 | * | 0.170 | ns | 1.000 | ns |
| | 14 | < 0.001 | *** | < 0.001 | *** | < 0.001 | *** |

**Supplementary Fig. 2c**

| Condition | Trial 1 | | Trial 2 | | Trial 3 | |
|---|---|---|---|---|---|---|
| | *p*-value | Significance | *p*-value | Significance | *p*-value | Significance |
| Myr-K 100 /Pep 1 – Myr-K 100/Pep 0.2 | 0.290 | ns | 0.592 | ns | 1.000 | ns |
| Myr-K 500/Pep 0.2 – Myr-K 100/Pep 0.2 | 1.000 | ns | 0.111 | ns | 1.000 | ns |
| Myr-K 500/Pep 1 – Myr-K 100/Pep 0.2 | 1.000 | ns | 0.977 | ns | 1.000 | ns |
| Myr-K 500/Pep 0.2 – Myr-K 100/Pep 1 | 0.180 | ns | 1.000 | ns | 1.000 | ns |
| Myr-K 500/Pep 1 – Myr-K 100/Pep 1 | 0.700 | ns | 0.288 | ns | 1.000 | ns |
| Myr-K 500/Pep 1 – Myr-K 500/Pep 0.2 | 1.000 | ns | 0.072 | ns | 1.000 | ns |

**Supplementary Fig. 5a**

| Trial | Condition | Mock | MeJA 1 µM | GR24 | GR24/ MeJA 0.1 µM | GR24/ MeJA 1 µM |
|---|---|---|---|---|---|---|
| 1 | MeJA 1 µM | 1.000 (ns) | - | - | - | - |
| | GR24 | 1.000 (ns) | 1.000 (ns) | - | - | - |
| | GR24/MeJA 0.1 µM | 1.000 (ns) | 1.000 (ns) | 1.000 (ns) | - | - |
| | GR24/MeJA 1 µM | 1.000 (ns) | 1.000 (ns) | 1.000 (ns) | 1.000 (ns) | - |
| | GR24/MeJA 10 µM | 1.000 (ns) | 1.000 (ns) | 1.000 (ns) | 1.000 (ns) | 1.000 (ns) |
| 2 | MeJA 1 µM | 1.000 (ns) | - | - | - | - |
| | GR24 | 1.000 (ns) | 1.000 (ns) | - | - | - |
| | GR24/MeJA 0.1 µM | 1.000 (ns) | 1.000 (ns) | 1.000 (ns) | - | - |
| | GR24/MeJA 1 µM | 1.000 (ns) | 1.000 (ns) | 1.000 (ns) | 1.000 (ns) | - |
| | GR24/MeJA 10 µM | 1.000 (ns) | 1.000 (ns) | 1.000 (ns) | 1.000 (ns) | 1.000 (ns) |
| 3 | MeJA 1 µM | 1.000 (ns) | - | - | - | - |
| | GR24 | < 0.001 (***) | 0.019 (*) | - | - | - |
| | GR24/MeJA 0.1 µM | < 0.001 (***) | 0.002 (**) | 1.000 (ns) | - | - |
| | GR24/MeJA 1 µM | < 0.001 (***) | 0.002 (**) | 1.000 (ns) | 1.000 (ns) | - |
| | GR24/MeJA 10 µM | < 0.001 (***) | 0.002 (**) | 1.000 (ns) | 1.000 (ns) | 1.000 (ns) |

**Fig. 3a**

| Condition | Trial 1 | | |
|---|---|---|---|
| | *p*-value | Hedge's *g* | Significance |
| Mock – MeJA 1 µM | 1.000 | 0.247 | ns |
| Mock – GR24 | 0.010 | 1.414 | ** |
| Mock – GR24/MeJA 0.1µM | 0.001 | 2.078 | *** |
| Mock – GR24/MeJA 1 µM | < 0.001 | 1.982 | *** |
| Mock – GR24/MeJA 10 µM | 0.001 | 2.692 | ** |
| MeJA 1 µM – GR24 | 0.024 | 1.237 | * |
| MeJA 1 µM – GR24/MeJA 0.1 µM | 0.001 | 1.963 | ** |
| MeJA 1 µM – GR24/MeJA 1 µM | < 0.001 | 1.894 | *** |
| MeJA 1 µM – GR24/MeJA 10 µM | 0.003 | 2.347 | ** |
| GR24/MeJA 0.1 µM – GR24 | 0.413 | 0.830 | ns |
| GR24/MeJA 1 µM – GR24 | 0.027 | 0.982 | * |
| GR24/MeJA 10 µM – GR24 | 1.000 | 0.559 | ns |
| GR24/MeJA 1 µM – GR24/MeJA0.1 µM | 1.000 | 0.209 | ns |
| GR24/MeJA 10 µM – GR24/MeJA 0.1µM | 1.000 | 0.430 | ns |
| GR24/MeJA 10 µM – GR24/MeJA 1 µM | 1.000 | 0.606 | ns |

**Supplementary Fig. 5b**

| Condition | Trial 2 | | | Trial 3 | | |
|---|---|---|---|---|---|---|
| | *p*-value | Hedge's *g* | Significance | *p*-value | Hedge's *g* | Significance |
| Mock – MeJA 1 µM | 1.000 | 0.417 | ns | 1.000 | 0.230 | ns |
| Mock – GR24 | < 0.001 | 1.305 | *** | < 0.001 | 1.581 | *** |
| Mock – GR24/MeJA 0.1 µM | < 0.001 | 1.835 | *** | < 0.001 | 3.270 | *** |
| Mock – GR24/MeJA 1 µM | < 0.001 | 1.844 | *** | < 0.001 | 2.279 | *** |
| Mock – GR24/MeJA 10 µM | 0.129 | 2.261 | ns | < 0.001 | 3.452 | *** |
| MeJA 1 µM – GR24 | 0.436 | 1.046 | ns | < 0.001 | 1.568 | *** |
| MeJA 1 µM – GR24/MeJA 0.1 µM | 0.048 | 1.330 | * | < 0.001 | 3.290 | *** |
| MeJA 1 µM – GR24/MeJA 1 µM | 0.003 | 1.677 | ** | < 0.001 | 2.292 | *** |
| MeJA 1 µM – GR24/MeJA 10 µM | 0.379 | 1.831 | ns | < 0.001 | 3.469 | *** |
| GR24/MeJA 0.1 µM – GR24 | 1.000 | 0.248 | ns | 0.006 | 1.825 | ** |
| GR24/MeJA 1 µM – GR24 | 0.130 | 1.003 | ns | 0.057 | 1.266 | ns |
| GR24/MeJA 10 µM – GR24 | 1.000 | 0.319 | ns | 0.036 | 1.662 | * |
| GR24/MeJA 1 µM – GR24/MeJA 0.1 µM | 1.000 | 1.212 | ns | 1.000 | 0.238 | ns |
| GR24/MeJA 10 µM – GR24/MeJA 0.1 µM | 1.000 | 0.737 | ns | 1.000 | 0.327 | ns |
| GR24/MeJA 10 µM – GR24/MeJA 1 µM | 1.000 | 0.830 | ns | 1.000 | 0.028 | ns |

**Fig. 3c**

| Condition | *p*-value | | | | | |
|---|---|---|---|---|---|---|
| | Trial 1 | | Trial 2 | | Trial 3 | |
| | T med | TG med | T med | TG med | T med | TG med |
| TG med | 0.250 (ns) | | 0.001 (**) | | 0.004 (**) | |
| TGM med | < 0.001 (***) | < 0.001 (***) | < 0.001 (***) | < 0.001 (***) | < 0.001 (***) | < 0.001 (***) |

**Supplementary Fig. 7b**

| Condition | Trial 1 | | Trial 2 | | Trial 3 | |
|---|---|---|---|---|---|---|
| | *p*-value | Significance | *p*-value | Significance | *p*-value | Significance |
| Subculture (Single) – Initial asymbiotic culture | 0.012 | * | 0.022 | * | < 0.001 | *** |
| Subculture (Block) – Initial asymbiotic culture | 0.763 | ns | 0.217 | ns | 0.789 | ns |
| Subculture (Single) – Subculture (Block) | 0.002 | ** | < 0.001 | *** | < 0.001 | *** |

**Fig. 3a**

| Condition | Trial 1 | | Trial 2 | | Trial 3 | |
|---|---|---|---|---|---|---|
| | *p*-value | Significance | *p*-value | Significance | *p*-value | Significance |
| Inoculated with AS – SS (2 WAI) | 0.343 | ns | 0.606 | ns | 1.000 | ns |
| Inoculated with AS – SS (4 WAI) | 0.343 | ns | 1.000 | ns | 1.000 | ns |
| Inoculated with AS – SS (6 WAI) | 0.343 | ns | 1.000 | ns | 1.000 | ns |
| Inoculated with AS – SS (8 WAI) | 0.343 | ns | 1.000 | ns | 1.000 | ns |

**Fig. 3b**

| Condition | Trial 1 | | Trial 2 | | Trial 3 | |
|---|---|---|---|---|---|---|
| | *p*-value | Significance | *p*-value | Significance | *p*-value | Significance |
| Inoculated with AS – SS (2 WAI) | < 0.001 | *** | 0.033 | * | 0.014 | * |
| Inoculated with AS – SS (4 WAI) | 0.009 | ** | 0.232 | ns | 0.231 | ns |
| Inoculated with AS – SS (6 WAI) | 0.045 | * | 0.232 | ns | 0.487 | ns |
| Inoculated with AS – SS (8 WAI) | 0.045 | * | 0.232 | ns | 0.487 | ns |

**Supplementary Fig. 9a**

| Condition | Trial 1 | | Trial 2 | | Trial 3 | |
|---|---|---|---|---|---|---|
| | *p*-value | Significance | *p*-value | Significance | *p*-value | Significance |
| Inoculated with AS – SS (2 WAI) | < 0.001 | *** | 0.038 | * | 0.001 | ** |
| Inoculated with AS – SS (4 WAI) | 0.014 | * | 0.232 | ns | 0.231 | ns |
| Inoculated with AS – SS (6 WAI) | 0.088 | ns | 0.232 | ns | 0.487 | ns |
| Inoculated with AS – SS (8 WAI) | 0.088 | ns | 0.232 | ns | 0.487 | ns |

**Fig. 4f**

| Trial | Condition | | *p*-value | Significance |
|---|---|---|---|---|
| 1 | Mock – | AS (200) | < 0.001 | *** |
| | | AS (400) | < 0.001 | *** |
| | | SS | < 0.001 | *** |
| | SS – | AS (200) | 0.729 | ns |
| | | AS (400) | 0.473 | ns |
| | AS (200) – | AS (200) | 0.971 | ns |
| 2 | Mock – | AS (200) | 0.398 | ns |
| | | AS (400) | 0.022 | * |
| | | SS | 0.004 | ** |
| | SS – | AS (200) | 0.065 | ns |
| | | AS (400) | 0.800 | ns |
| | AS (200) – | AS (200) | 0.311 | ns |
| 3 | Mock – | AS (200) | < 0.001 | *** |
| | | AS (400) | < 0.001 | *** |
| | | SS | < 0.001 | *** |
| | SS – | AS (200) | 0.009 | ** |
| | | AS (400) | 0.045 | * |
| | AS (200) – | AS (200) | 0.793 | ns |

**Supplementary Table 4.** Composition of the modified Long Ashton medium. After mixing all compounds, pH was adjusted to 6.8 by adding NaOH.

| Chemicals | Final concentration |
|---|---|
| $Ca(NO_3) \cdot 4H_2O$ | 354 mg $L^{-1}$ |
| KCl | 54 mg $L^{-1}$ |
| $KH_2PO_4$ | 2.7 mg $L^{-1}$ |
| Fe(III)-EDTA | 42 mg $L^{-1}$ |
| $MgSO_4 \cdot 7H_2O$ | 185 mg $L^{-1}$ |
| KI | 1.2 mg $L^{-1}$ |
| $MnCl_2 \cdot 4H_2O$ | 14 mg $L^{-1}$ |
| $H_3BO_3$ | 22 mg $L^{-1}$ |
| $ZnCl_2$ | 1.7 mg $L^{-1}$ |
| $CuCl_2 \cdot 2H_2O$ | 0.47 mg $L^{-1}$ |
| $CoCl_2 \cdot 6H_2O$ | 0.17 mg $L^{-1}$ |
| $Na_2MoO_4 \cdot 2H_2O$ | 1.4 mg $L^{-1}$ |
| $KNO_3$ | 101 mg $L^{-1}$ |
| PIPES (pH7.5) | 305 mg $L^{-1}$ |
